# Evolution of the Metazoan Protein Domain Toolkit Revealed by a Birth-Death-Gain Model

**DOI:** 10.1101/2025.07.03.659011

**Authors:** Yuting Xiao, Maureen Stolzer, Larry Wasserman, Dannie Durand

## Abstract

Domains, sequence fragments that encode protein modules with a distinct structure and function, are the basic building blocks of proteins. The set of domains encoded in the genome serves as the functional toolkit of the species. Here, we use a phylogenetic Birth-Death-Gain model to investigate the evolution of this protein toolkit in metazoa. Given a species tree and the set of protein domain families in each present-day species, this approach estimates the most likely rates of domain origination, duplication and loss.

Statistical hierarchical clustering of domain family rates reveals sets of domains with similar rate profiles, consistent with groups of domains evolving in concert. Moreover, we find that domains with similar functions tend to have similar rate profiles. Interestingly, domains with functions associated with metazoan innovations, including immune response, cell adhesion, tissue repair, and signal transduction, tend to have the fastest rates.

We further infer the expected ancestral domain content and the history of domain family gains, losses, expansions, and contractions on each branch of the species tree. In contrast to recent reports of widespread loss during metazoan evolution, we observe little evidence of genome streamlining. Rather, our analysis reveals an ongoing process of domain family replacement and resizing, consistent with extensive remodeling of the protein domain repertoire. The use of a powerful, probabilistic Birth-Death-Gain model reveals an unexpected level of genomic plasticity and a striking harmony between the evolution of domain usage in metazoan proteins and organismal innovation.

## 1 Introduction

The fascinating diversity of forms observed in the animal kingdom is a driving force in metazoan biology. Metazoan innovations include complex multicellularity, with spatiotemporal cell type differentiation; cell junctions; organs; radial and bilateral body plans; endocrine signaling; innate and adaptive immune response; muscles, bones, cartilage, and a complex nervous system (DeSalle and Schierwater, 2007; Boero et al., 2007; Camara et al., 2007; Deline et al., 2018). The expanding wealth of genome sequences from metazoa and their closest unicellular relatives is driving a growing body of work investigating the relationship between the evolution of the metazoan protein-coding repertoire and innovations on other levels of biological organization. Phylogenetic comparative methods are being used to probe the genetic basis of Metazoan innovation broadly (Wang et al., 2012; Technau and Schwaiger, 2015; Paps and Holland, 2018; Fernández and Gabaldón, 2020; Guijarro-Clarke et al., 2020; Zmasek and Godzik, 2011, 2012; Domazet-Lošo et al., 2024; Suga et al., 2013; Grau-Bové et al., 2017; López-Escardó et al., 2019; Richter et al., 2018), or with a focus on specific systems, including the bilaterian body plan (Heger et al., 2020), the metazoan nervous system (Emes et al., 2008; Ryan and Grant, 2009; Liebeskind et al., 2015, 2016), and membrane proteins (Nam et al., 2015; Attwood et al., 2017).

Ancestral state reconstruction proceeds from a set of phylogenetic profiles, one for each family, representing the number of copies of the family encoded in each present day genome. Protein domains, domain combinations, and full-length proteins provide three related, but distinct views of the protein repertoire encoded in a genome (Fig. 1). Several lines of evidence suggest that the emergence of novel multidomain architectures, via exon shuffling and other mechanisms, played an important role in early metazoan evolution (Patthy, 2021; King et al., 2008). While most studies of the protein coding repertoire have focused on full length protein families, domains (Zmasek and Godzik, 2011; Moore and Bornberg-Bauer, 2012; Wang et al., 2012; López-Escardó et al., 2019; Suga et al., 2013) and domain combinations (Grau-Bové et al., 2017; Zmasek and Godzik, 2012) have also been considered.

**Fig. 1:**
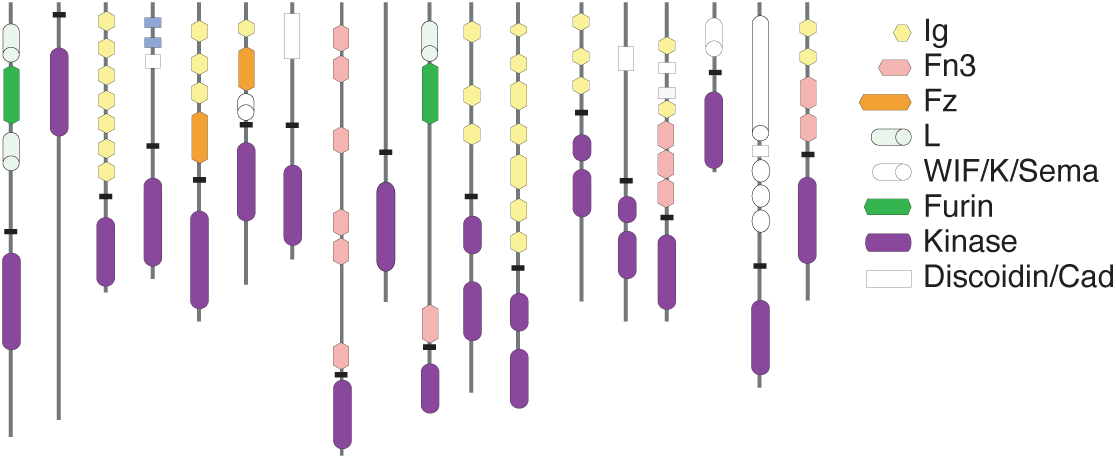
Domain architectures in a complex multidomain protein family, the receptor subfamily of the human protein tyrosine kinases, adapted from (Robinson et al., 2000).

Phylogenetic comparative methods infer the ancestral protein coding repertoire and state changes along the branches of the species tree that best explain the present-day phylogenetic profiles with respect to a set of assumptions about evolutionary processes. Parsimony methods infer ancestral states that minimize the number of gains and losses (Swofford and Maddison, 1987). Dollo parsimony, which is widely used in comparative genomic studies, applies the further restriction that a character may be lost multiple time, but only be gained once (Farris, 1977). Probabilistic methods that do not minimize the number of inferred changes can also be used (Harmon, 2019; Pagel, 1999). These methods implement a parameterized mathematical model that embodies a set of assumptions about the evolutionary changes that can occur and the probability of those changes.

Here, we apply a probabilistic, birth-death-gain (BDG) model, implemented in the Count software package (Csűrös, 2010; Csűrös and Miklós, 2009), to investigate the evolutionary processes that gave rise to the protein domain repertoire encoded in the genomes of 21 species spanning the evolutionary trajectory of the metazoa and their closest unicellular relatives (Fig. 2). In Count, the evolutionary events that modify the protein coding repertoire, birth (i.e., duplication of a domain already encoded in the genome), death (loss of a domain), and gain (*de novo* domain acquisition), are explicit parameters of the model. Given a species tree and present-day domain counts, Count infers birth, death, and gain rates that maximize the likelihood of the present day repertoire, as well as the probability of ancestral states and state changes on the tree. The inferred domain rates are not only a means to obtain an ancestral reconstruction, but also a source of biological insight.

**Fig. 2:**
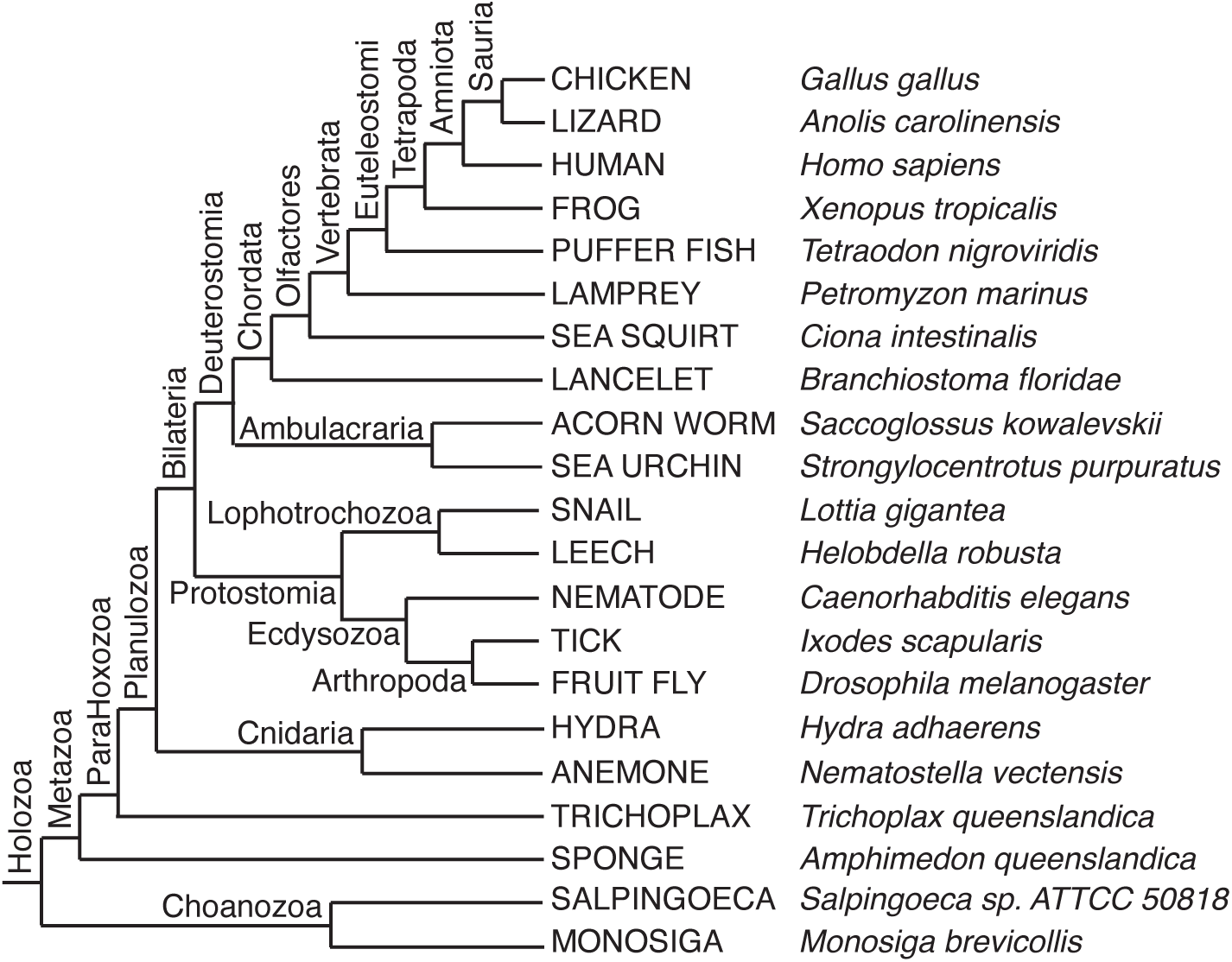
Species tree of the 21 metazoan species used in our analysis, adapted from Philippe et al. (2009)

As a probabilistic model, BDG does not exert a preference for minimal change. Further, there is no *a priori* preference for losses versus gains. The inclusion of a gain event in the model allows for multiple independent gains of the same domain family in different branches of the species tree. Ancestral reconstruction using a stochastic model infers expected behavior reflecting contributions of many possible evolutionary trajectories. This flexible model has the potential to reveal plasticity, if such exists, that could not be uncovered with a more constrained model. We investigate the evolution of the protein domain repertoire through the lens of these model properties.

## 2 Methods

### 2.1 Data

The Birth-Death-Gain analysis was carried out using domain data from the genomes of 21 holozoan species (19 metazoa and 2 choanoflagellates) and three outgroups: *Neurospora crassa*, *Dictyostelium discoideum*, and *Arabidopsis thaliana*.

Genomes used in this analysis were chosen to probe major metazoan clades, while also considering genome quality. In addition, taxa were selected to avoid very short or very long branches along the backbone of the phylogeny to minimize artifacts and convergence problems associated with extreme branch lengths. For this reason, ctenophores are not represented in this study.

The species phylogeny used in this study (Fig A1) is based on that of Philippe et al. (2009). Early metazoan branching order is controversial. In the phylogeny used in this work, sponge and Planulozoa are sister taxa, where the planulozoan clade comprises Cnidaria and Bilateria (Lartillot and Philippe, 2008; Philippe et al., 2009; Telford et al., 2015). While the sister clade to sponge is often referred to as Eumetazoa, we do not use this terminology because the term “Eumetazoa” implies a clade that includes ctenophores, which are not represented in our data set. Recently, the placement of Trichoplax with respect to Cnidaria and Bilateria has been called into question, with some analyses placing Trichoplax and Cnidaria as sister taxa (Simakov et al., 2020; Laumer et al., 2018). However, other recent work (Laumer et al., 2019; Simion et al., 2017; Najle et al., 2023) continues to support the topology used here (Figs. 2 and A1).

Domain annotations in these 24 genomes were obtained from the SUPERFAMILY database (Oates et al., 2015), which classifies protein domains based on structural and evolutionary relationships using the SCOP (Structural Classification of Proteins) framework (Andreeva et al., 2020; Pandurangan et al., 2019). The SUPERFAMILY classification is hierarchical, with domain models corresponding to SCOP families and SCOP superfamilies. Members of the same superfamily share a structural core, but sequence similarity within the superfamily may be low. We used these top-level assignments for all domain families in our dataset. Since only one level of the hierarchy was used, we use the term “domain family” (and not “superfamily”) throughout this manuscript.

For each of the 24 genomes, domain annotations of protein-coding sequences corresponding to the longest transcript per gene were downloaded from the SUPERFAMILY database, version 1.75 (Oates et al., 2015; Pandurangan et al., 2019). Domain family counts for each species were then tabulated from this data, resulting in 1,434 distinct families. The resulting table served as input for the Birth-Death-Gain model described below.

For functional analyses in this study, we use the domain-specific functional ontology (Vogel et al., 2004, 2005) provied by the SUPERFAMILY database (https://supfam. org/SUPERFAMILY/function.html). The “InterPro2GO” project provides manual annotation of domains with GO terms, based on experimental characterizations of the domain’s function (Burge et al., 2012; Blum et al., 2025). However, at the time of writing, only 30% (395) of the 1,283 families in this study have been assigned one or more GO terms. Of the 1,283 domain families in our study, 1,220 are annotated in the Vogel ontology. For functional enrichment analysis, we considered only the 1,092 domain families annotated with informative labels or identified as viral proteins.

### 2.2 Ancestral reconstruction

#### 2.2.1 Phylogenetic Birth-Death-Gain (BDG) model

We investigated the evolution of domain family sizes using the phylogenetic BDG model implemented in the Count software package (Csűrös, 2010). The model takes as input a species tree, *T* = (*V, L, B*), with nodes *V*, leaves *L* ⊂ *V*, and branches *B*, and a matrix of present-day family sizes. Each family is represented as a vector of length |*L*|, where entry *S* contains the number of instances of the family encoded in the genome of species *S* ∈ *L*. Count models family size evolution as a continuous-time Markov process, with transition probabilities that depend on both branch-specific and family-specific parameters. On branch *b* ∈ *B*, family size increases with probability (*κ* + *λn*)*t_b_* and decreases with probability *µnt_b_*, where *n* is the family size, *t_b_* is the length of *b*, and *λ*, *κ*, and *µ* are the rates of birth, gain, and death events, respectively. The probability that a domain duplication will occur increases with the number of domains in the family, whilst the probability of a gain is independent of family size. A birth-death model (*κ* = 0) entails the implicit assumption that all families in present-day species were present in their common ancestor. A non-zero gain rate allows for origination of novel domain families in any lineage of the species tree. Family sizes in the common ancestor (i.e., the root of *T*) are assumed to follow a Poisson distribution with mean, *ϕ*.

The Count model permits rate variation across species tree lineages, input families, or both. Branch-specific variation allows for different values of the branch length (*t_b_*) and the birth and gain rates, *λ_b_* and *κ_b_*, for each branch *b* ∈ *B*. The loss rate, *µ_b_*, is set to 1 by normalization. Parameter variation across families is modeled as a discretized gamma distribution with *C* categories, where the value for each category is the mean of the corresponding quantile. Each parameter value for family *f* is represented as a weighted sum over the category values for that parameter, where the weights are estimated from prior probabilities as discussed in greater detail, below. When using the full model, the birth, death, and gain rates are the product of a branch-specific rate, a family-specific rate, and a family-specific scaling factor, *σ_f_* . Thus, the birth rate for family *f* on branch *b* is *λ* = *σ_f_ λ_f_ λ_b_*. The death and gain rates are defined analogously.

In Count, parameter estimation is implemented as an iterative numerical maximization procedure (Csűrös and Miklós, 2009). Parameter estimation is considered to have reached convergence when the increase in the log-likelihood between consecutive iterations is less than 0.01. This inference process estimates the Poisson parameter (*ϕ*) and the branch length parameters (*t_b_, λ_b_, κ_b_,* ∀*b* ∈ *B*). The optimization process also fits the discretized gamma distributions for family-specific variation The Count manual (Csűrös, 2009) recommends that the maximum likelihood parameter estimation procedure be carried out in stages of increasing model complexity to facilitate convergence. In the first stage, the values of *σ, κ, λ, µ* and *ϕ* are estimated with no rate variation. These serve as starting estimates for the next stage, which models branch-specific variation, only. The resulting values are used as starting values for the third stage, which comprises both branch- and family-specific variation.

The results of this maximum likelihood inference procedure are used to calculate the expected rates and ancestral sizes of each family, *f*, in a separate procedure provided by Count. In this second pass, for each parameter *π* ∈ {*σ_f_, κ_f_, λ_f_, µ_f_* }, the expected value of *π* is the weighted sum ^L^ *p_f_* (*c, π*)·*π_c_*, where *p_f_* (*c, π*) is the probability that *π_f_* is in category *c*.

Ancestral states are also calculated in the second pass. To reduce the number of parameters, Count only considers three states: the family is absent (*n* = 0), a singleton (*n* = 1), or multi-member (*n >* 1). For each ancestral node in *v* ∈ *V*, the probabilities *p_f_* (*v, n* = 0), *p_f_* (*v, n* = 1), and *p_f_* (*v, n >* 1) are determined for every family *f* . The expected numbers of families that are absent, singletons and multimers in ancestor *v* are given by ^L^ *p_f_* (*v, n* = 0), *_f_ p_f_* (*v, n* = 1) and ^L^ *p_f_* (*v, n >* 1), respectively. Count also outputs the probability of each possible state change (0 to 1, 1 to many, many to 1, 1 to 0) on every *b* ∈ *B*.

#### 2.2.2 Phylogenetic Birth-Death-Gain (BDG) inference

We applied the Count model to the species tree in Figure A1 and phylogenetic profiles described in Data above. Twenty-two extremely large families (Table A2) could not be processed by Count and were excluded from further analysis. A table of domain counts in the remaining 1,412 families served as the input to the phylogenetic Birth-Death-Gain analysis. Guided by prior empirical evaluation of the impact of model choice on Count performance (Stolzer et al., 2014), in this study we used both branch-specific variation and family-specific variation with *C* = 2 categories (i.e., a fast rate and a slow rate), for all four parameters {*σ_f_, κ_f_, λ_f_, µ_f_* }.

The maximum likelihood parameter estimation procedure was carried out in stages, as recommended by the Count manual (Csűrös, 2009). The inferred parameter values obtained in the final stage are given in Table A3 and Fig A2. From these, the expected family-specific rates (Fig A3) and ancestral states were determined. The three outgroup species were included in this analysis for the purposes of rate inference only. After removing domain families specific to these outgroups, 1,283 domain families remained. Scaled rates, *σ_f_ λ_f_*, *σ_f_ κ_f_*, and *σ_f_ µ_f_*, were calculated for the 1,283 holozoan domain families (Fig A4) and used in downstream analyses.

#### 2.2.3 Dollo parsimony inference

We also performed ancestral reconstruction using Dollo parsimony to allow for comparison with prior studies. The Dollo parsimony method infers ancestral states under the assumption that a domain family can be gained only once in a lineage, but can be lost multiple times. Ancestral states were inferred using the Dollo parsimony implementation in Count, with the domain family count table for all 24 species and the full species tree as inputs.

### 2.3 Definitions

#### Core and non-core domain families

For a given ancestral node, we define core domain families as those present in all present-day descendants of that ancestor. In this study, we focus on the data set core (i.e., families in all 24 genomes), as well as the holozoan core, metazoan core, bilaterian core, and euteleost core families for comparison and discussion (Table A5).

#### Domain family repertoire

In present-day genomes, the domain family repertoire refers to number of copies encoded in the genome of each of the 1,283 holozoan domain families. In ancestral genomes, the domain repertoire refers to the expected number of families encoded in the genome as estimated by Count, as described above. The ancestral domain repertoire can further be partitioned into the expected numbers of singleton families and multimer families. It is possible to estimate the number of families, but not the number of domains, encoded in an ancestral genome, because Count does not estimate the expected number of copies in a multimer family. Although the actual present-day family sizes are known, in some figures we represent present-day genomes in terms of the fraction of families that are singletons or multimers, to allow for comparison between ancestral and present-day domain family repertoires.

#### Domain events

In the context of domain family evolution, birth and death events correspond to domain duplication and deletion, respectively. Here, we use the term duplication broadly to comprise any evolutionary process that creates a new copy of a domain somewhere in the genome. This includes both gene duplication and domain duplication. Similarly, domain deletion can result from the loss of a domain or the loss of an entire gene.

#### Domain family events

For each domain family, *f*, and each branch *b* = (*u, v*) in the species tree, Count estimates the probabilities of the following *family events*: *f originated* on *b* (i.e., *f* was not encoded in species *u* and was encoded in species *v*, where *u* is ancestral to *v*); *f expanded* on *b* (i.e., was encoded in a single copy in *u* and more than one copy in *v*); *f contracted* on *b* (i.e., was encoded in more than one copy in *u* and a single copy in *v*), *f* was *extinguished* on *b* (i.e., *f* was encoded in *u* but not in *v*); or *f* remained unchanged. Note that domain family events are not synonymous with domain events. For example, a family extinction is always the result of a domain loss, but a domain loss only results in a family extinction, when the family is size one.

For each family, Count estimates the probability of a given event occurring on a given branch. In this study, we say family *f* was gained in species *v* if the difference between the origination and extinction probabilities of *f* on *b* = (*u, v*) is greater than 0.6. Similarly, we say *f* was lost on *b* if this difference is less than -0.6.

The family event probabilities are used to estimate the expected number of state changes of each type on each branch. The expected number of families that originated on *b* is the sum of the probability, for each family *f*, that *f* originated on *b*. The expected numbers of families that expanded, contracted or expired on *b* are calculated analogously.

#### Family Turnover

For a given branch, *b* = (*u, v*), comparing the expected repertoires of *u* and *v* gives the net change in the numbers of families encoded. The number of family events can greatly exceed change in the size of the repertoires. Family turnover refers to situations where the sum of the expected family gains and expected family losses on *b* substantially exceeds the net change in families encoded in *v*, relative to *u*.

### 2.4 Clustering of event rates

We applied Statistical Hierarchical Clustering (SHC) to the 1,283 family-specific scaled rate profiles (*σ_f_ λ_f_*, *σ_f_ κ_f_*, and *σ_f_ µ_f_*) to determine whether groups of domains share distinct rate profiles. SHC (Kimes et al., 2017) extends traditional hierarchical clustering by providing a statistical assessment for the clustering results obtained. Hierarchical clustering is the process of repeatedly merging clusters of points, generating progressively larger clusters until a termination criterion is satisfied. At each iteration, the pair of clusters to be merged is selected according to a protocol that depends on the inter-cluster distances between all pairs. Inter-cluster distances are recalculated and the process repeats. Hierarchical clustering methods differ in the metric used to calculate pairwise distances, the protocol for selecting a pair to merge, and the termination criterion.

SHC offers a choice of metrics and merging protocols and a termination criterion that ensures that the resulting clustering deviates significantly from a null model. At every merge point in the agglomerative procedure, SHC calculates a test statistic that compares pairwise distances within each cluster, considered separately, with pairwise distances in the set resulting from their union. The statistic under the null hypothesis is estimated by Monte Carlo sampling from a Gaussian distribution derived from the clusters to be merged. The p-value is the fraction of replicates that had a better statistic than the genuine bipartition. The SHC termination condition ensures that the significance of each cluster in the final clustering exceeds a user-provided significance threshold, *α*, after family-wise error correction. SHC guarantees *p < α* for every cluster in the clustering; in some cases, the p-values associated with the individual clusters are much smaller than *α*. We report the largest (i.e., least significant) p-value for each clustering.

The scaled rates were clustered with four metrics (Euclidean, Pearson, Manhattan, Maximum) and five agglomeration strategies (Average, Median, Complete, Centroid, Single) available in the SHC package. Prior to clustering with SHC, the data was centered and scaled using the R Scales package (Wickham et al., 2025). Domain families with identical rate profiles were removed, reducing the input by 64 families. For downstream analyses, the excluded domain families were restored by assigning each family to the same cluster as their corresponding identical rate profile. Of these 20 combinations, 10 yielded significant clusterings at the *α* = 0.05 level (Table A1). At this level of significance, the chance expectation of a significant clustering is one in 20. All clusterings have a mean silhouette (Rousseeuw, 1987; Maechler et al., 2023) above 0.5, suggesting that the clusterings are reasonably coherent.

We next asked whether the 10 significant clusterings capture similar relationships between domain families, using Normalized Mutual Information (NMI), an information-theoretic measure that quantifies the amount of shared information between two distributions. The raw mutual information is normalized by the clustering entropy, providing a scale-independent measure that ranges from zero to 1. NMI is widely used in computational biology due to its interpretability and sensitivity to clustering granularity (Vinh et al., 2010; Strehl and Ghosh, 2002; Cover, 1999; Rodriguez et al., 2019; Chiquet et al., 2020). The NMI values for our clusterings suggest that, with the exception of clusterings that used average linkage, there is substantial overlap in the information they encode (Fig. A5).

## 3 Results

We investigate the evolution of the protein toolkit in 21 genomes, comprising two choanoflagellate genomes and 19 metazoan genomes, ranging from sponge to man. These genomes encode 1,283 domain families, as defined by the SUPERFAMILY database; 501 core families are found in all 21 genomes and 547 are common to metazoa. Among the non-core families, 84 are species-specific (Table A4). Notably, 37 of these are found in sponge. The present-day phylogenetic profiles of non-core families show additional taxonomic structure (Fig. A6), including families found uniquely in choanoflagellate, metazoan, and vertebrate genomes. Idiosyncratic absences are also observed. A substantial number of families that are broadly represented in metazoan genomes are absent from the lamprey genome, and to a lesser extent from the sea squirt, nematode, tick, and fruitfly genomes..

### 3.1 Family-specific domain rate variation

We used the Birth-Death-Gain model implemented in Count (Csűrös, 2010) to infer the event rates that maximize the likelihood of present-day holozoan domain family sizes. For each family, *f*, among the 1,283 distinct domain families in the 21 species analyzed, this yielded family-specific gain, duplication and loss rates (*λ_f_, µ_f_* and *κ_f_*), as well as a scaling factor (*σ_f_*) that captures the rate of change of the family, overall.

To examine the relationship between domain rates and domain function, we used a manually curated, domain-specific ontology (Vogel et al., 2004, 2005), consisting of 50 detailed functions grouped into 7 general functions (Table A6). The ontology is constructed such that each domain is mapped to at most one general and one detailed function category. In our dataset, 63 domain families lack annotations because they had not yet been identified at the time of curation; 148 domain families were not assigned a function or were annotated as “unknown function” or “General”.

Figure 3 shows the distribution of the overall scaling factor (*σ_f_*) and the scaled duplication (*σ_f_ λ_f_*), gain (*σ_f_ κ_f_*), and loss (*σ_f_ µ_f_*) rates in each functional category represented in our data set. Functional categories are ordered according to the median scaling factor (*σ_f_*, bottom panel) in each category. The distribution of the median scaling factor across functional categories is reminiscent of a sigmoid function, with a rapid transition from slow to fast rates. Many functions with low medians are associated with basic cellular processes of all cells across the tree of life, including information processing functions (e.g., transcription, translation, DNA replication), transport and carbon metabolism. At the high end, functional categories like cell adhesion and immune response are associated with the development of multicellularity and metazoan immune system.

**Fig. 3:**
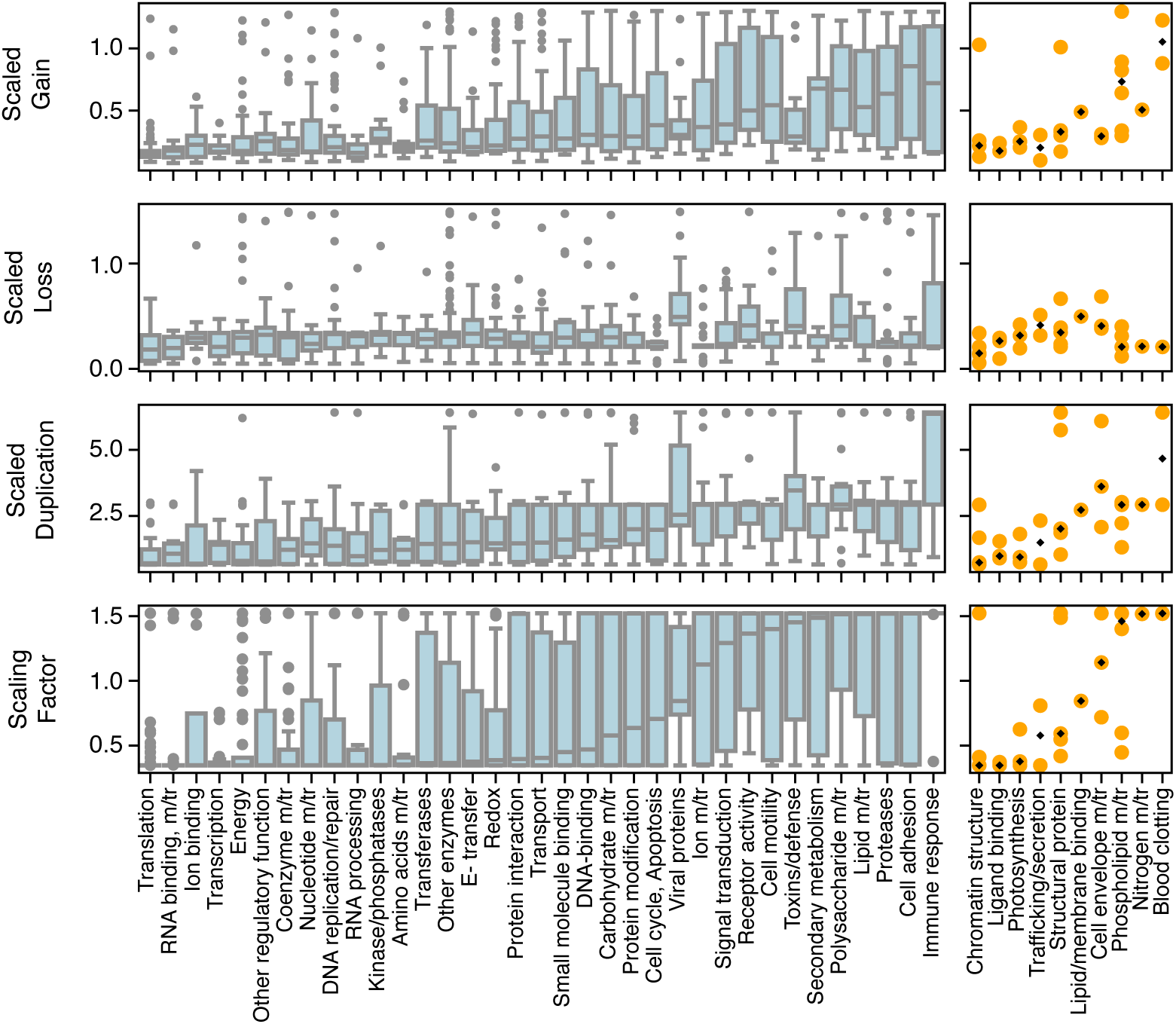
Box-and-whisker plots showing the scaling factor (*σ_f_*) and scaled duplication (*λ_f_*), loss (*µ_f_*), and gain (*κ_f_*) rate distributions of 1072 domain families distributed across functional categories (63 recently discovered domain families lack annotations). Categories are ordered by ascending median scaling factor value. Rates in categories with fewer than 10 families are plotted as individual points, shown at right. Note that m/tr stands for metabolism and transport.

### 3.2 Ancestral domain content

We next probed how the domain family repertoire varies across the species tree. The fraction of the 1,283 families encoded in present-day genomes ranges from 66% in the choanoflagellate *M. brevicollis* to 81% in human; 41% to 61% of these families, respectively, are present in more than one copy. Within this range, protein domain content shows considerable variation. The domain repertoire in Nematostella and sponge is noticeably larger than the repertoires of their closest relatives. Within Bilateria, the domain complement is, on average, somewhat smaller in protostomes than in deuterostomes, with some notable exceptions. The lamprey genome, for example, encodes fewer domain families than any other genome in this data set. To better understand the history of this domain repertoire variation, we used Count to estimate the expected number of families that are present, as singletons or multimers, in each ancestral genome (Fig. 4b). According to this reconstruction, approximately 68% of the 1,283 present-day domain families were encoded in the holozoan ancestor; slightly more than half of those families were multicopy. The fraction of families in ancestral genomes along the Bilateria/Deuterostome backbone grows gradually to 78% in the bilaterian ancestor. The expected domain famaily repertoires encoded in the protostome and lophotrochozoan ancestors are similar in size to that of Bilateria. In contrast, the repertoires and, especially, the multicopy repertoires in Ecdysozoa and Athropoda are somewhat smaller.

**Fig. 4:**
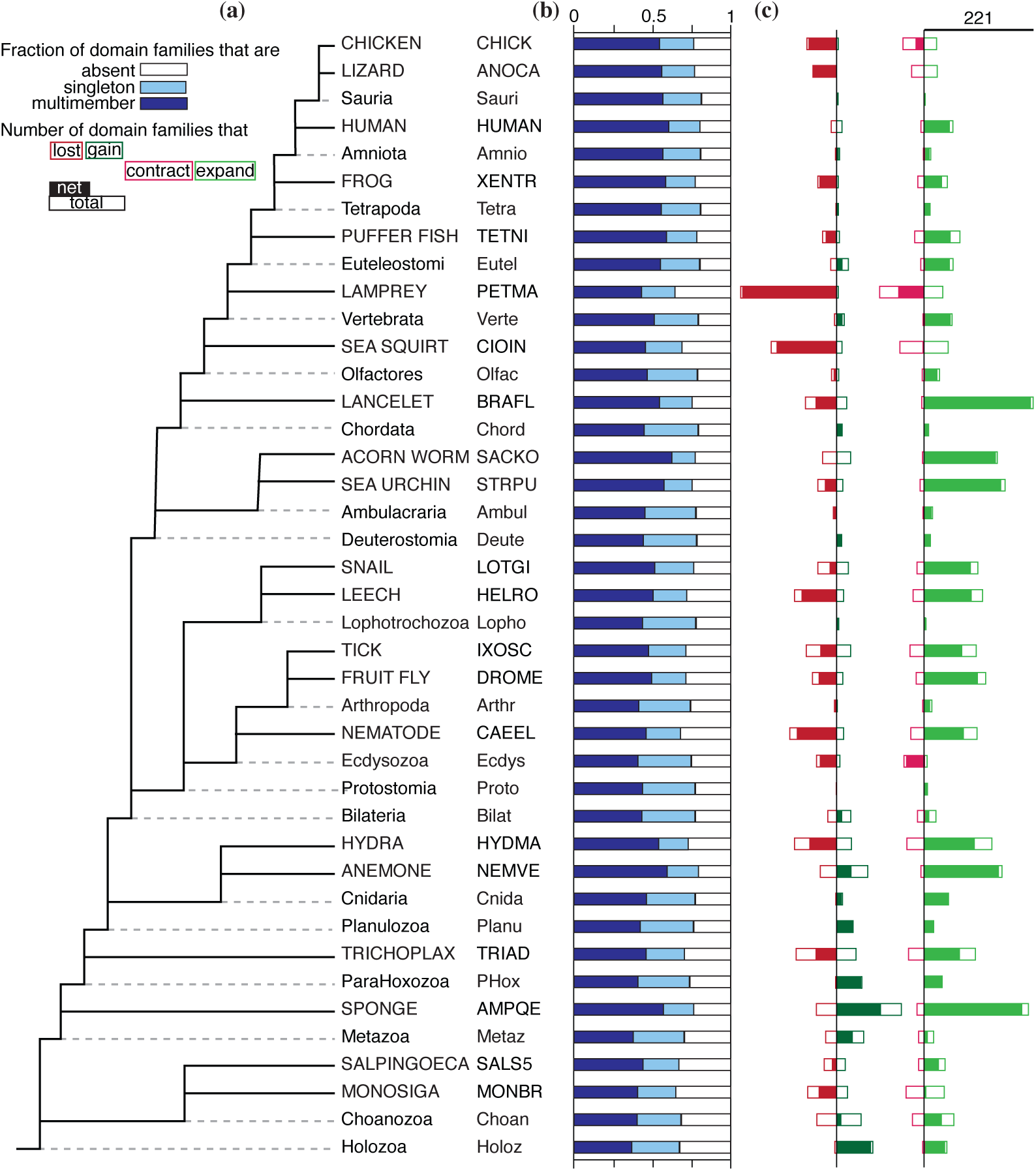
**(a)** Phylogeny of the holozoan genomes used in this study. **(b)** The expected fraction of the 1,283 domain families that are absent (white), singleton (light blue), and multimer families (dark blue) in present-day and ancestral species. **(c)** The expected numbers of families that originated and were extinguished, shown in outline in dark The expected fractions of families that expanded and contracted, shown in outline in light green and pink. The net change is shown as a solid bar.

This reconstruction provides an estimate of the size of the protein domain repertoire encoded in ancestral genomes, but does not reveal to what extent the specific families that made up these repertoires changed over the course of metazoan evolution. To assess the plasticity of the ancestral repertoire, we calculated, for each branch in the species tree the expected number family gains, expansions, contractions, and losses on that branch (Fig. 4c). Many branches, including the lineages leading to ParaHoxozoa, Planulozoa, Cnidaria, Protostomia, Deuterostomia, Chordata, and Amniota, exhibit modest family gain and expansion, which is consistent with the gradual increase in the number and size of domain families in the ancestral backbone species.

In other lineages, however, the inferred family events reveal much more activity than is apparent from the changes in the size of the ancestral domain repertoire. For example, the modest increase in the number of families on lineage leading to the Metazoan ancestor is the result of substantial family gains offset loss of other families, resulting in a small net increase. The specific families that made up the multicopy repertoire also changed substantially. Although many families changed in size, the increase in the number of multicopy families is quite modest, because the number of family expansions is only slightly larger than the number of family contractions, A similar pattern of family turnover, with substantial actual change, but little net change, is seen in the Bilaterian, Choanozoan and Euteleost lineages, as well as in several terminal branches. Rather than pure growth, this pattern suggests a process of domain repertoire remodeling, where some families grow, other families shrink, and still other families are replaced with new ones.

We also see lineages, including many terminal lineages, where family losses exceed family gains, resulting in a net loss in the number of domain families encoded in the genome. In some species (e.g., lancelet and sea urchin), this is accompanied by substantial number of family expansions. In some cases, such as trichoplax and hydra, modest family turnover is superimposed on this broad pattern of specialization. In other species, notably Ecdysozoa and lamprey, both family loss and family contraction dominate.

Overall, in metazoan lineages, the evolution of the protein domain repertoire can be characterized by four evolutionary modes (Fig. 5):

- Genome expansion, in which both the number and the size of domain families increases.
- Genome remodeling, where net change in the number of families belies a much greater degree of family origination and extinction. In addition, expansion in some families is offset by contraction in other families, resulting in a relatively small change in the number of domain copies (from any family) encoded in the genome.
- Specialization, in which the number of domain families decreases, but those families that are retained increase in size. The variety of tools is decreased - monly those types that are needed for the specialized task are retained.
- Genome streamlining, in which both the number and the size of the families in the genome are reduced.

**Fig. 5:**
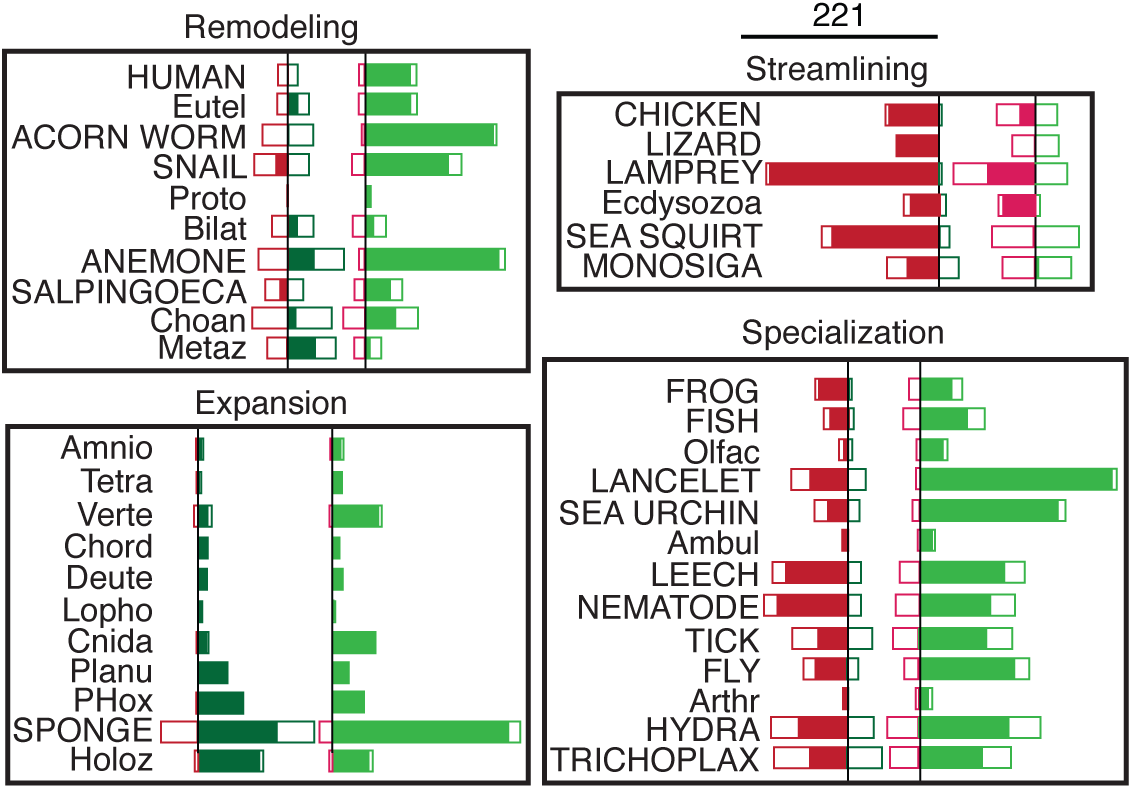
Strategies of domain repertoire evolution.

### 3.3 Are groups of domains evolving in concert?

The emergence of phenotypic novelty is associated with expansion and elaboration of existing cellular pathways and protein complexes, as well as the appearance of new ones. This hypothesis predicts that domains that mediate protein-protein interactions in the same cellular pathways and proteins complexes will tend to have similar rate profiles. In addition, the frequency of domain pairs (domain bigrams) in multidomain sequences deviates significantly from expectation, given domain unigram frequencies (Cui et al., 2024, 2022; Vogel et al., 2005), suggesting that the formation of domain combinations is selectively constrained. Under these constraints, domains that frequently co-occur will also tend to have similar duplication, gain, and loss rates.

With this in mind, we asked whether there are groups of domains that exhibit greater similarity in rate profiles than expected by chance. To identify such groups, if they exist, we applied statistical hierarchical clustering (Kimes et al., 2017) to the set of 1,283 family-specific scaled rate profiles {*σ_f_ λ_f_, σ_f_ µ_f_*, *σ_f_ κ_f_* }, using five agglomeration strategies and four metrics, as described in Methods. Ten of these 20 combinations yielded significant clusterings at the *α* = 0.05 level (Table A1). Since less than one clustering in 20 is expected by chance at this significance level, this is strong evidence that groups of domains share evolutionary trajectories.

We selected the Euclidean-Centroid clustering for further analysis, because it is highly significant (*p* = 3e−17) and is representative of the majority of clusterings obtained (Fig. A5). Each cluster in this grouping has a characteristic rate profile (Fig. 6). Cluster I, by far the largest cluster, has slow gain, duplication and loss rates. Clusters II and II, which are intermediate in size, have moderate duplication rates and high gain rates, but differ in their loss rates. The three smallest clusters (IV - VI) are characterized by very high duplication rates.

**Fig. 6:**
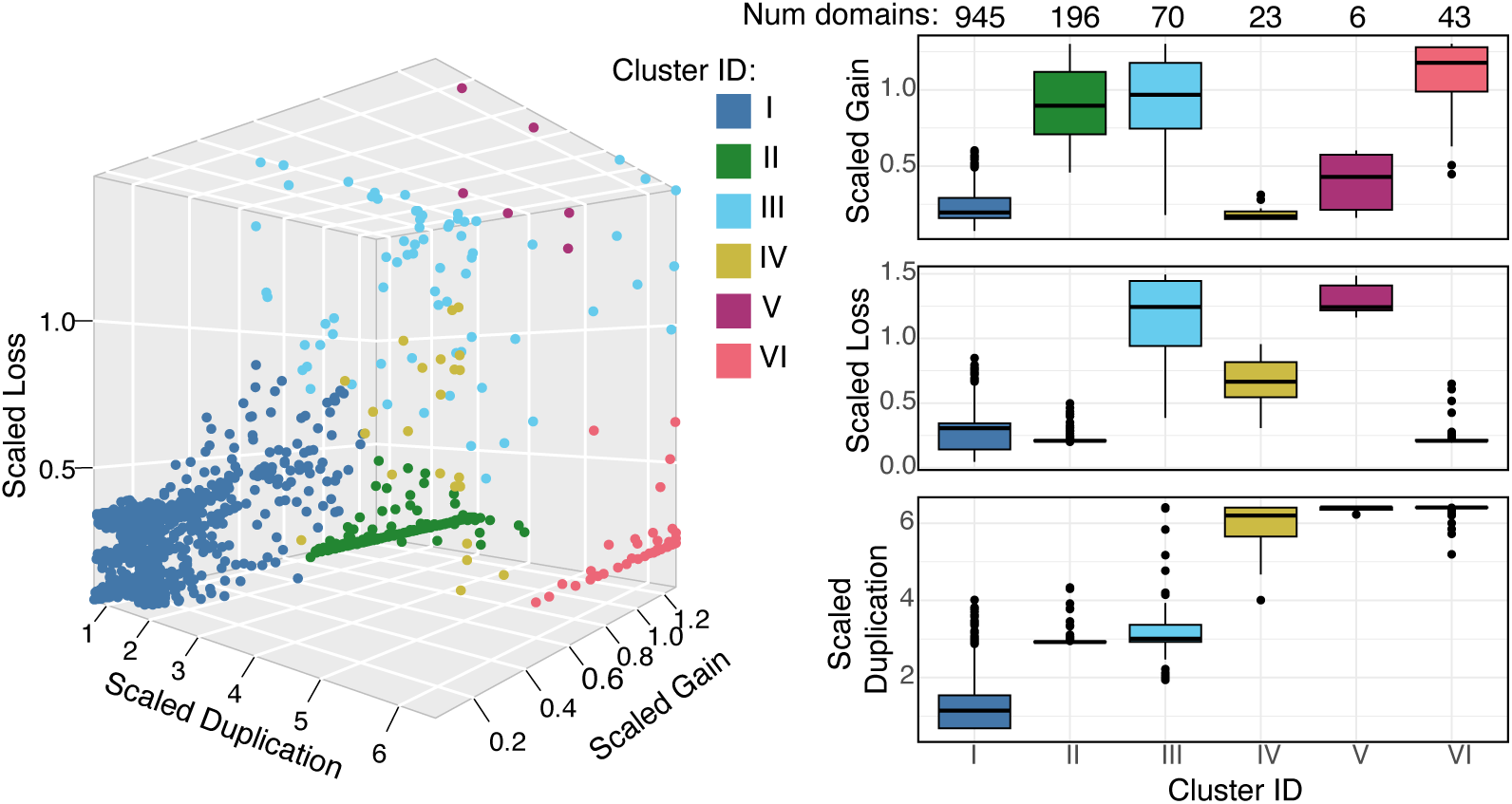
**(a)** Scaled gain, loss, and duplication rates, clustered using centroid linkage with the Euclidean metric. Dot colors indicate cluster assignments. **(b)**Scaled gain, loss and duplication rate distributions, by cluster. Clusters are ordered by median duplication rate.

In order to better interpret these differentiated rate profiles, we examined the functional profile (see Methods) for each cluster (Tables 1, A7 and A8). Since Cluster IV - VI are very small and share a very high duplication rate, we combined them for the purposes of functional enrichment analysis. After multiple testing correction, we find that Cluster I is significantly enriched for domains with information processing functions (hypergeometric test, *p <* 0.05). Domains with roles in extra- and intra-cellular processes are underrepresented. In contrast, in Cluster II, information processing is underrepresented and intra-cellular processes are over represented. Taken together, domain families with extra-cellular functions are enriched in the three clusters with very high duplications. Only one domain with an information processing function is found in these three clusters, combined.

**Table 1:** Distribution of General Functions across scaled rate clusters. Bold: *p <* 0.05 with multiple testing correction. p-values (hypergeometric test) are given in Table A9.

| General Function | I |  | II |  | III |  | IV-VI |  | Total |  |
| --- | --- | --- | --- | --- | --- | --- | --- | --- | --- | --- |
| Extra-cellular | 16 | 2% | 13 | 7% | 4 | 7% | 13 | 22% | 46 | 4% |
| General | 69 | 9% | 17 | 9% | 6 | 10% | 4 | 7% | 96 | 9% |
| Information | 157 | 20% | 10 | 5% | 3 | 5% | 1 | 2% | 171 | 16% |
| Intra-cellular | 99 | 12% | 54 | 29% | 8 | 13% | 14 | 24% | 175 | 16% |
| Metabolism | 324 | 41% | 57 | 30% | 31 | 51% | 13 | 22% | 425 | 39% |
| Regulation | 110 | 14% | 36 | 19% | 7 | 11% | 9 | 16% | 162 | 15% |
| Viral | 11 | 1% | 0 | 0% | 2 | 3% | 4 | 7% | 17 | 2% |
| **Total** | 786 |  | 187 |  | 61 |  | 58 |  | 1092 |  |

Next, we asked how the domains in each cluster are distributed across present-day genomes (Fig. 7). About a third of the 945 domains in Cluster I are present in all 21 genomes in this data set. Another 10% are present in a single species, consistent with lineage specific gains. The remaining families in Cluster I have a patchy distribution. Perhaps a quarter of domain families in this cluster are present in a one or a small number of genomes only. This includes the 37 sponge-specific domain families, for example. Clusters II and VI are primarily made up of core families (Table A5). Almost 75% of Cluster II families and 40% of Cluster VI families are encoded in all present-day holozoan genomes in our study. Eighty percent of Cluster VI families are in the Bilaterian core. The vast majority of domain families in both clusters are multicopy. In contrast, the taxonomic distribution of Cluster III is patchy: 68 of 70 families are encoded in at least two, but not all, genomes. Cluster is similarly patchy. Cluster IV families also have sparse phylogenetic profiles, but with less patchiness: half of Cluster IV families are present in only one species and another quarter are specific to a single taxonomic clade.

**Fig. 7:**
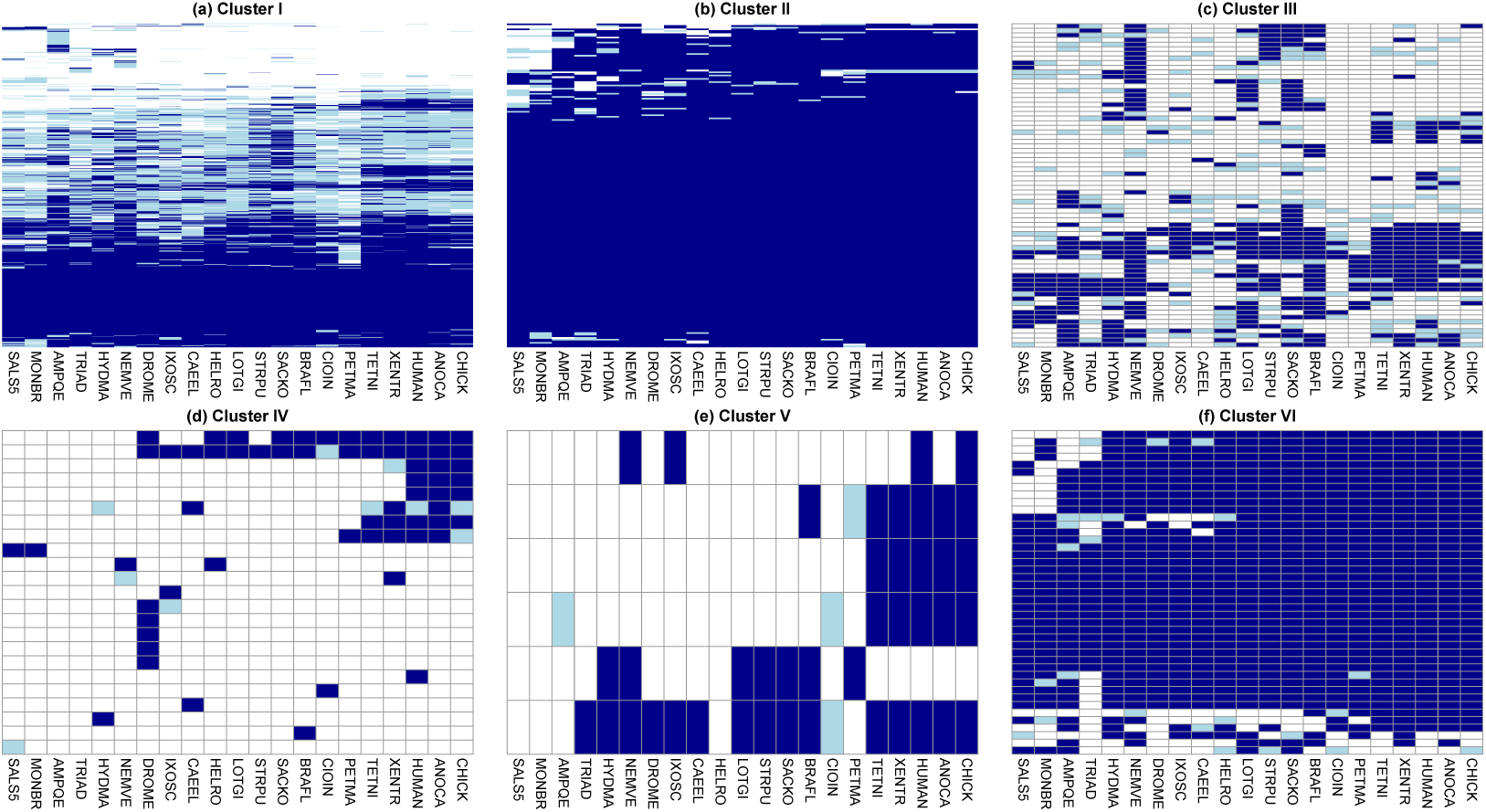
Distribution of domain families in each cluster over 21 present-day, holozoan genomes (dark blue: present in two or more copies; light blue present in one copy; white: absent). Dendogram of domain family phylogenetic profiles in each cluster generated using the pheatmap package in R (Kolde, 2018).

The taxonomic distributions of individual clusters range from very dense to patchy to very sparse. The patchy distributions must be the result of parallel gains, parallel losses or reversals. Clusters with very dense or very sparse clusters can be explained with minimal parallelism, but might also conceal a higher degree of homoplasy. To better understand the evolutionary processes that gave rise to these present-day distributions, we examined the ancestral states (Fig. 8) and the pattern of family gain, expansion, contraction and loss associated with the families in each clusters (Fig. 9).

**Fig. 8:**
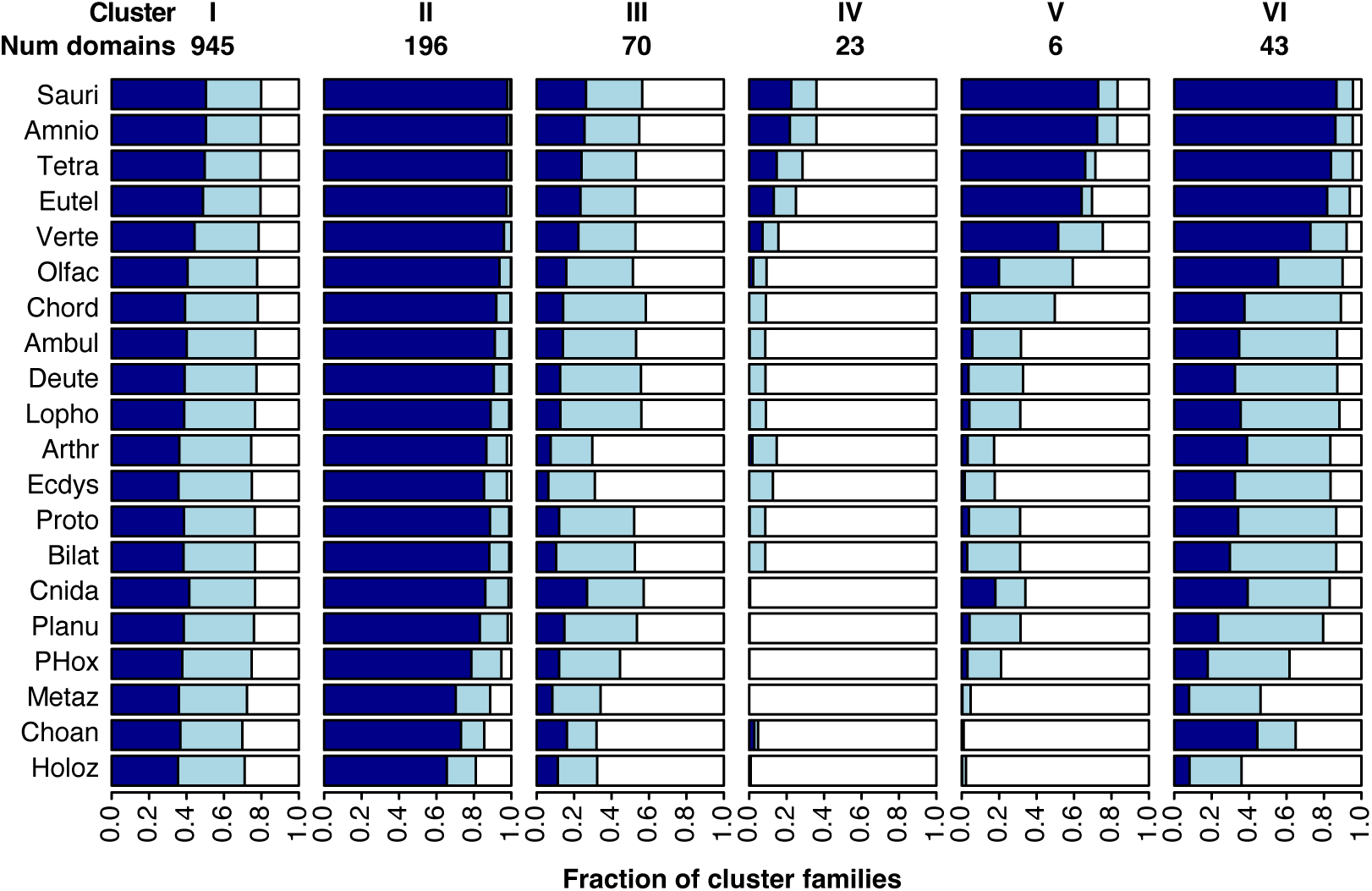
Ancestral family sizes and family events in Clusters I-VI. **(a)** The expected frequency of absent (white), singleton (light blue), and multimer families (dark blue) in ancestral species in each cluster. Family counts are normalized by cluster size. **(b)** The expected fraction of families that originated, expanded, contracted and were lost in non-terminal lineages.

**Fig. 9:**
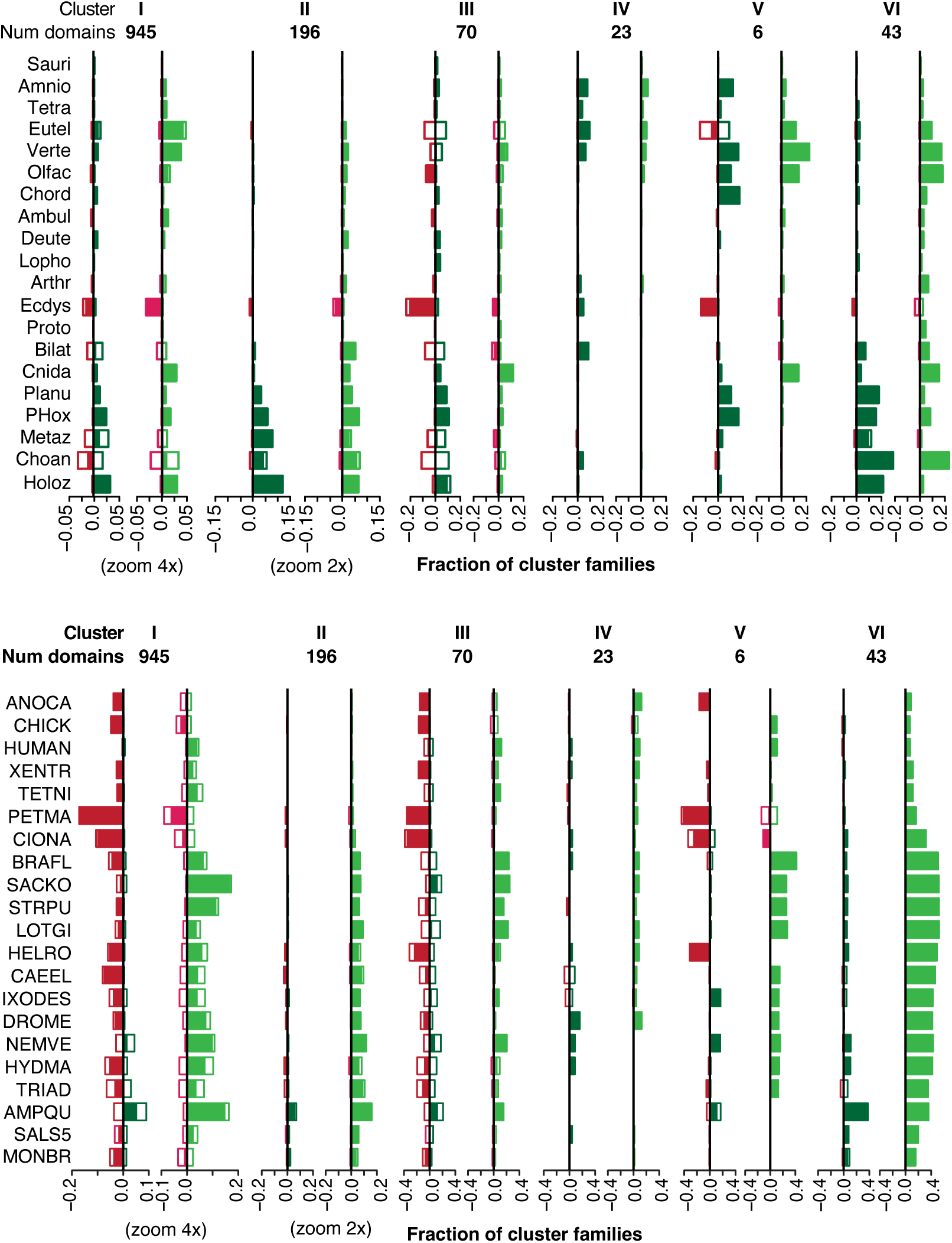
Expected family events in Clusters I-VI. The expected fraction of families that originated, expanded, contracted and were lost in ancestral (top panel) and terminal (bottom panel) lineages.

Cluster I families display a mix of core, patchy and sparse phylogenetic profiles. This mix is consistent with the relatively broad event rate distributions in Cluster I. The median gain, duplication, and loss rates are low, but the upper quantiles in all three distributions are in the mid-range. In the reconstruction, each ancestral genome encodes an expected 75% to 80% of Cluster I families, with multimember families ranging from 40% in the holozoan ancestor to 50% in euteleosts. The frequencies of singleton and multicopy Cluster I families encoded in ancestral genomes are remarkably stable. However, the expected number of family gain and family loss events are consistent with ongoing family turnover (Fig. 9), which suggests that the actual families vary from ancestor to ancestor.

The histories of family events in Clusters I and III show family turnover in the choanozoan, metazoan and bilaterian ancestors. Similarly, turnover is evident in the euteleost ancestor in Clusters III and V. Many terminal lineages also indicate substantial turnover, especially in Cluster III. In Cluster I, family events on terminal branches indicate a mixture of turnover and specialization. In all three clusters, the present-day patchy distributions, combined with ongoing loss, replacement and resizing of families, is suggestive of extensive remodeling of the protein domain repertoire.

In contrast, very little turnover was inferred for Cluster IV families. The present-day profiles of this cluster are so sparse that this cluster can be explained by a small number of gains.

It is instructive to compare their evolutionary histories of Clusters II and IV, which have similar, dense taxonomic distributions. The reconstruction of domain families encoded in ancestral genomes suggests that roughly 80% of Cluster II families were already present in the holozoan ancestor; virtually all families were encoded in the planulozoan ancestor. In all ancestral nodes, the majority of families in the expected repertoire are encoded in two or more copies. The expected family events in ancestral lineages consist of family gains and expansions (Fig. 9). In summary, Cluster II exhibits a consistent, gradual expansion of a stable Metazoan protein domain toolbox.

In contrast to Cluster II, the acquisition of the full complement of Cluster VI multicopy families occurred more recently than their common ancestor. Although an expected 85% of families were encoded in the Bilaterian ancestor, most of these were singletons; increases in copy number occurred much later in the Chordate clade. The distribution of inferred family gains across ancestral species is similar to that of Cluster II. However, family expansion continued in recent ancestors and in many invertebrate terminal lineages (Fig. 9), consistent with the higher duplication rates in Cluster VI. Note that this inferred history differs from a parsimony-based ancestral reconstruction, which would predict a full suite of multicopy families in the Bilaterian ancestor.

### 3.4 Parallel Gains and Losses

The present-day phylogenetic distribution (Fig. 7) reveals a number of families that are present in distantly related taxa, but absent in their closest relatives, which suggests a history of independent gains, independent losses, and reversals. These patchy distributions are particularly evident in clusters with high loss and/or gain rates. Moreover, the expected family events (Fig. 9) are consistent with parallel expansions in terminal lineages.

To determine whether individual families are sustaining parallel gains and parallel losses, for each family, we examined the probability of a family gain or loss event on each branch of the species tree (Fig. A7 - A10). We consider that a family sustained a high-confidence gain (respectively, loss) event if the difference between the gain and loss probabilities is greater than 0.6 (respectively, less than -0.6). This analysis identified 115 families that have a history of at least two high-confidence gain events. Many of these family sustained multiple independent gains, especially in Clusters III and VI which have high gain rates. Taken together, these 115 families comprise 315 parallel gains. Many instances of families with a high probability of extinction in distinct lineages were also identified: 300 families exhibited a history of two or more high-confidence loss events, representing 1,085 parallel family loss events, in total.

These trends are exemplified by the six families in Cluster V (Fig. 10). In the BDG reconstruction, three of the six families (47266, 69349, 82671) sustained parallel loss events. These histories are consistent with the Dollo parsimony criterion. Two more families (57392, 117773) have histories with multiple, independent gains. Note that for both families, a much larger number of events would be required for a history that satisfies the Dollo parsimony criterion. The BDG reconstruction for the remaining family (54452) does not satisfy any parsimony criterion: this family is gained in the vertebrate ancestor, but is immediately lost in lamprey. A history comprising a gain in the euteleost ancestor requires fewer events.

**Fig. 10:**
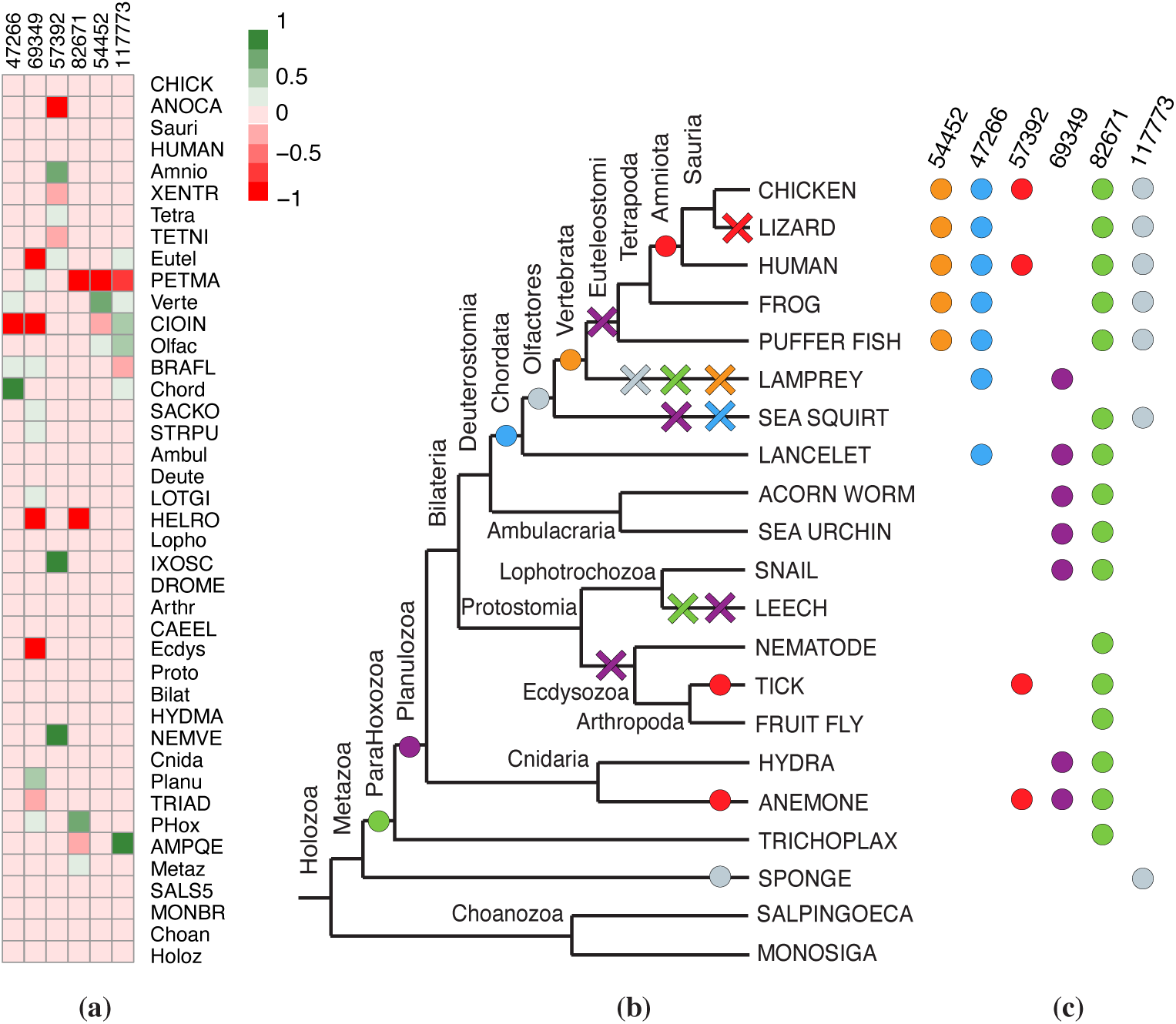
Family gain and loss probabilities for the six domain families in Cluster V. **(a)** Probability of family gain minus probability of family loss on each species tree branch. **(b)** High-confidence family gain and loss events are depicted as circles and crosses, respectively, on the species tree. **(c)** Family presence in present-day genomes is indicated by colored circles. Families are labeled with their SUPERFAMILY IDs.

The prevalence of parallel gains inferred by the BDG model led us to wonder how Dollo parsimony, which is widely used for investigating the evolution of the protein-coding repertoire (Zmasek and Godzik, 2011; Fernández and Gabaldón, 2020; Guijarro-Clarke et al., 2020; Domazet-Lošo et al., 2024; Fairclough et al., 2013; Qiu et al., 2015, 2018; Bowles et al., 2020, e.g.), would perform on this data set. According to the Dollo parsimony criterion, a family can be gained only once, but can be lost multiple times. Unlike the BDG method, which allows for parallel events of either type, under the Dollo criterion, a patchy distribution can only be explained by an early gain, followed by multiple independent losses. To investigate the impact of this constraint, we applied Dollo parsimony reconstruction method implemented in Count (Csűrös, 2010) to infer the number of families gained and lost on each branch, using the same input data as for the BDG model. We examined the net change (gains minus losses) estimated by both models on each branch (Fig. 11). The results show that Dollo parsimony has an overwhelming preference for family losses, compared with the BDG model. In the Dollo reconstruction, family losses exceeded gains on 13 of 20 internal branches in the species tree; in the BDG reconstruction, losses dominated gains on only 5 of 20 branches. Considering all branches combined, Dollo inferred a total of 300 gain events and 2,239 loss events, while the BDG model inferred 895 expected family gains and 1,477 expected losses.

**Fig. 11:**
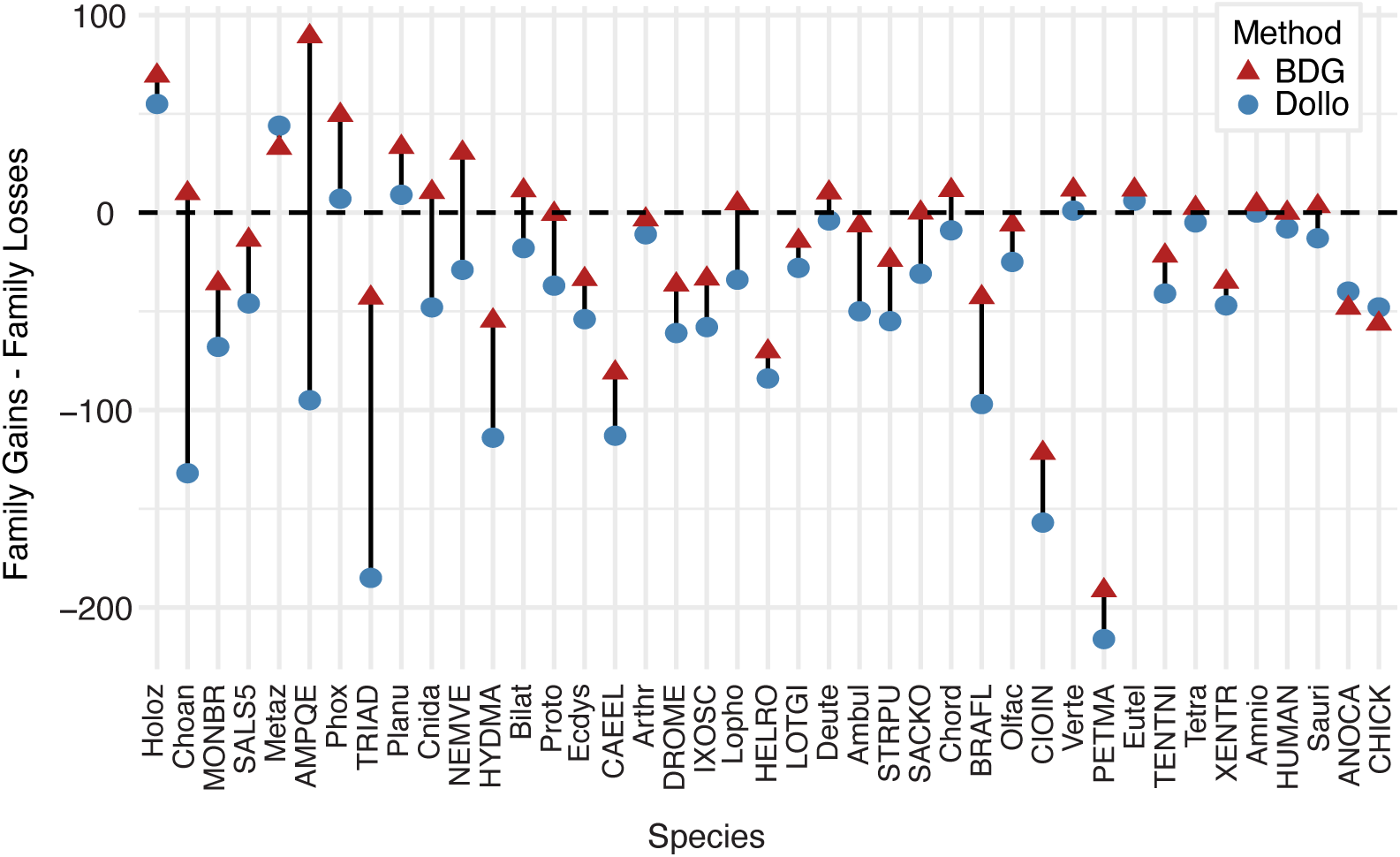
Net change in family representation on each species tree lineage. Expected family gains minus expected family losses, as estimated with the Birth-Death-Gain model (red triangles). Gains minus losses estimated with Dollo parsimony (blue circles).

## 4 Discussion

In this work, we investigated the evolution of the protein domain repertoire in 21 holozoan species using the Birth-Death-Gain model implemented in Count (Csűrös, 2010; Csűrös and Miklós, 2009). We used the full Count model, which includes both branch- and family-specific rate variation. Incorporating family-specific variation has been shown to improve model fit, yielding more accurate estimators without sacrificing generality (Stolzer et al., 2014). Importantly, family-specific rates not only improve accuracy, but also provide insight into the evolutionary trajectories of individual domain families.

Using Count, we inferred the rates of domain gain, duplication and loss. When domain event rates are examined in light of functional properties, our results reveal that domains associated with basic cellular processes (e.g., transcription, translation, DNA replication, metabolism) tend to have slower evolutionary rates. In contrast, domain families with functions related to innovations that appear during metazoan evolution (e.g., cell adhesion, immune response, receptor activity and signal transduction) exhibit elevated rates.

We further asked whether there are cohorts of domains that are evolving in concert. We posit that if major events in organismal evolution are linked to changes in protein-mediated cellular processes, then domains associated with the requisite pathways and protein complexes will have similar rate profiles. Indeed, statistical hierarchical clustering (Kimes et al., 2017) of domain family rates revealed clusters that exhibit statistically robust rate differentiation. These clusters also share functional and historical attributes, consistent with a link between protein innovation and the emergence of phenotypic novelties.

Using the same model, we estimated the family sizes in ancestral genomes, as well as the expected family gains, expansions, contractions and extinctions in each lineage. We observed four distinct evolutionary modes that shape the protein domain repertoire in metazoa. A small number of lineages are characterized by genome streamlining, that is, net loss of domain families and reduction in size among those that remain. Specialization of the protein domain repertoire, where the loss of some families is offset by a size increase in others, is a much more common evolutionary mode. A third set of lineages is characterized by gradual growth and expansion. Finally, and most compelling, we observe lineages where family turnover dominates. Notably, this process of remodeling, where some families are lost and are replaced by others, was inferred in the ancestors of several major clades, including Metazoa, Choanozoa, Bilateria and Protostomia, that are associated with dramatic and highly successful phenotypic innovations.

The limited importance of genome streamlining in this reconstruction stands in stark contrast to recent studies that report widespread loss in Metazoa (Guijarro-Clarke et al., 2020; Domazet-Lošo et al., 2024) and other multicellular groups (Qiu et al., 2018; Bowles et al., 2020; Zmasek and Godzik, 2011). Many of these studies are based on analyses using Dollo parsimony, which, by design, resolves homoplasy with parallel losses. Reanalysis of our data set with Dollo parsimony dramatically increases the number of losses inferred. The ratio of losses to gains increases almost fivefold, from 1.7 losses for every gain obtained with the BDG model to 7.5 losses per gain as inferred by Dollo parsimony.

Our results demonstrate that, for this data set, the number of losses inferred is much greater with Dollo parsimony than with a Birth-Death-Gain model. While this discrepancy is noteworthy, the true history of the metazoan domain repertoire is unknown. We cannot rule out the possibility that Dollo parsimony is correctly inferring a history of massive loss. Nevertheless, there are several reasons to doubt that scenario. First, Dollo parsimony is designed to infer losses. Second, BDG models have the flexibility to model both parallel gains and parallel losses and, indeed, accounts of massive loss inferred with a BDG model have been reported (Csűrös and Miklós, 2009). In a data set truly characterized by parallel loss, we might expect the two methods to produce similar results. Finally, a recent study of the evolution of the eukaryotic protein domain repertoire also reported that Dollo parsimony obtained much higher losses than a maximum likelihood method (G’alvez-Morante et al., 2024). That study differed from ours in the taxonomic sample, the source of protein domain data, the implementation of Dollo parsimony and the specific probabilistic model used, emphasizing the potential role of Dollo parsimony in overestimating losses.

## Limitations and Future Perspectives

Ancestral reconstruction can be sensitive to the choice of taxa represented in the analysis. The species sample in this study is relatively small. Ctenophores were deliberately excluded to avoid convergence associated with very short branches. Xenacoelomorphans are also not represented in our data set. In addition, the number of genomes of unicellular holozoa and Teretosporea that have been sequenced has increased substantially (Ruiz-Trillo et al., 2023). Incorporating these in a future study would give a better assessment of the pre-metazoan state.

Uncertainty in the species tree can contribute to downstream uncertainty. Early metazoan branching order is controversial (Najle et al., 2023; Whelan et al., 2017; Giribet and Edgecombe, 2020; Laumer et al., 2018, 2019; Philippe et al., 2011; Simion et al., 2017; Dunn et al., 2008; Telford et al., 2015; Schultz et al., 2023; Simakov et al., 2020; Pandey and Braun, 2020; Redmond and McLysaght, 2021; Lartillot and Philippe, 2008; Philippe et al., 2009). One issue is the placement of ctenophores relative to porifera. This uncertainty does not directly impact our results because ctenophores are not represented in this study. The placement of Trichoplax with respect to Cnidaria and Bilateria has also been called into question, with some analyses placing Trichoplax and Cnidaria as sister taxa (Simakov et al., 2020; Laumer et al., 2018). However, other recent work (Laumer et al., 2019; Simion et al., 2017; Najle et al., 2023) continues to support the topology used here (Figs. 2 and A1).

Missing protein domain annotations, genome assembly and gene prediction errors, are another source of noise. Proteins contain domains that have not yet been characterized pose an even more fundamental problem. Fortunately, advances in AI-based protein structure prediction have dramatically increased both the size and the accuracy of protein domain resources (Paysan-Lafosse et al., 2025; Lau et al., 2024), offering exciting prospects for future studies.

Currently, there is no comprehensive functional annotation system for protein domains and this negatively impacted the depth of our functional analysis. Several projects currently underway will enable more comprehensive analyses of domain function in the not too distant future. The InterPro2GO project is a growing resource of manual annotation of domains with GO terms, based on experimental characterizations of the domain’s function (Burge et al., 2012; Blum et al., 2025). Complementing this effort, methodology to support automated mapping of Gene Ontology (GO) terms from proteins to domains is an active area of research (Weiner et al., 2008; Buchan and Jones, 2020; Fang and Gough, 2013; López and Pazos, 2013; Ulusoy and Dŏgan, 2024). More and better domain annotations will not only lead to more accurate reconstruction, but will also allow for more robust functional analyses. This analysis focuses on the protein domain repertoire, defined to be the set of domains encoded in the genome without regard to how the domains are distributed across protein coding genes. Since protein domains are basic units of protein function, the set of protein domains encoded in a genome constitutes the functional toolkit available for building complex functions in that species. It is also important to study how the evolution of domain architectures has contributed to metazoan evolution. Domain architectures are larger, more complex and more prevalent in animals than in other branches of the Tree of Life (Tordai et al., 2005). Many families that are integral to complex multicellularity consist of multidomain proteins with complex and varied architectures (Patthy, 2021). Very few studies have tackled phylogenetic comparative analyses of domain combinations in Metazoa (Zmasek and Godzik, 2012; Grau-Bové et al., 2017). This is an exciting, important and challenging area for future work.

Probabilistic methods suffer from disadvantages including model violations and overfitting. Despite its flexibility, the Birth-Death-Gain model is challenged by very long running times, which depend on the computational cost of a single iteration and the number of iterations required to reach convergence. Count’s likelihood maximization procedure requires summing over many latent variables. It is not uncommon for latent variable models to have poorly defined, multimodal likelihood functions which do not promote rapid convergence. It is difficult to predict how running times will scale as problem instances increase in size because the speed of convergence depends on the smoothness of the likelihood function in unexpected ways. Improvements in taxon sampling and domain architecture annotations are exciting directions for further study. At the same time, these will increase the scale of the problem. Whether BDG models can realistically be used to investigate these larger problems is an open question.

Ongoing sequencing efforts are changing our understanding of the Metazoa-specific gene complement. Genes that are associated with metazoan phenotypes and were previously believed to be specific to metazoa are increasingly found to have orthologs in close relatives to Metazoa, indicating that these families are were already present in pre-metazoan ancestors. This suggests that the genesis of novel phenotypes may lie not in the acquisition of new genes, but in the co-option of existing genes to new functional roles. This raises questions about the utility of ancestral reconstruction of the protein repertoire, not just for this study but for all studies that use this approach. Changes that led to the phenotypic innovation seen in Metazoa may be due to regulatory or epigenetic changes, for example.

## Methodological challenges to principled ancestral reconstruction

Phylogenetic comparative analysis provides a powerful framework for investigating the genetic underpinnings of phenotypic change. Ancestral reconstruction of the genomic features can reveal the historical coincidence of genetic and phenotypic shifts, predict functional links, identify potential convergent evolution and generate testable hypotheses. Phylostratigraphic analyses associate estimates of gene age with biological properties of interest. Present day phyletic distributions have contributed to evidence for *de novo* gene origination. However, this compelling framework entails substantial methodological challenges.

First, relating changes in gene family composition to phenotypic evolution requires correct identification of corresponding genetic features in different genomes. Comparative analyses can be compromised by biased or sparse taxon sampling, gene prediction errors, bias in orthology databases, and failure to recognize orthologs due to remote homology (Natsidis et al., 2021; Moyers and Zhang, 2015, 2017; Jain et al., 2019). This can be a problem for domains as well as genes (Kress et al., 2023; Nagy et al., 2011; Tassia et al., 2021).

Second, the assumptions underlying the ancestral reconstruction method must be aligned with the properties of the data set, including taxonomic and genomic characteristics and the idiosyncrasies of the underlying data. Dollo is just one of several parsimony methods that infer ancestral character states that minimize the number of state transitions, but differ in the relative importance of parallel gains, parallel losses, and reversals in resolving parsimony (Swofford and Maddison, 1987). Probabilistic models of character evolution also embody specific assumptions about character evolution (Harmon, 2019; Pagel, 1999), although the dependence on model specifics may be underappreciated. Some accounts simply report that ancestral states were inferred using “maximum likelihood” and give the name of a software package, but do not provide a clear statement of the underlying model and its assumptions.

Phylogenetic reconciliation (Stolzer, 2012), and related gene tree methods (e.g., Fernández and Gabaldón, 2020), are less widely used, but highly informative approach to inferring gene events and ancestral gene content. Parsimony-based reconciliation methods are computationally relatively efficient, yet are much less sensitive to the limitations of the parsimony assumption because the inference process is constrained by the information encoded in the gene tree topology. Phylogenetic reconciliation models can accommodate lateral gene transfer, allowing for multiple independent origins, albeit with an increase in computational complexity.

Whether parsimony or probability based, the quality of reconstruction will depend heavily on the extent to which the underlying assumptions of the model capture the properties of the biological system. Dollo parsimony is the algorithmic implementation of Dollo’s law of irreversibility, which posits that complex traits cannot be lost and regained in the same form. Whether these assumptions are always justified for complex traits is a long standing subject of debate (Elmer and Clobert, 2025). In contrast, in genomic studies, the appropriateness of the Dollo model (or any other model) is less frequently discussed.

In the context of genomic features, Dollo’s law of irreversibility implies that a genomic feature cannot be gained more than once (Farris, 1977). This assumption also underlies the phylostratigraphic approach to estimating gene age (Domazet-Lošo et al., 2007), which posits that the progenitor of a protein family arose in its cenancestor. When genetic features can be acquired horizontally, via lateral gene transfer (LGT) or introgression, for example, reconstruction paradigms that allow for multiple independent gains are required. This is particularly important in the face of increasing evidence for LGT in eukaryotes (Keeling, 2024; Sibbald et al., 2020; Van Etten and Bhattacharya, 2020). Dollo, which forbids multiple independent gains, tends to overestimate the both the age of a gene and the gene losses that occurred in the presence of horizontal gene flow (Capra et al., 2013).

In this study, analysis of the same data set with different ancestral reconstruction algorithms resulted in very different conclusions. With Dollo, domain loss is the dominant force: Losses exceed gains on two thirds of internal branches. With BDG, domain gain is the dominant force: Gains exceed losses on 75% of internal branches. This stark disagreement highlights the importance of selecting an ancestral reconstruction method with a compatible evolutionary model. Our results add to a growing body of work that suggests that the widespread use of phylostratigraphy and Dollo parsimony may be contributing to the perception that gene loss is a widespread phenomenon.

## Conclusions

Comparative phylogenetic analysis with a Birth-Death-Gain model reveals extensive and continual remodeling of the metazoan protein domain repertoire, suggesting an unexpected degree of plasticity. This is one of the few models capable of inferring event rates, providing a fresh perspective on the evolution of metazoan protein toolkit in relation to organismal innovation.

Evolution by genome streamlining, characterized by a dramatic increase in the ancestral genomic repertoire followed by genome reduction in multiple lineages, has been proposed as an important mode of genome evolution (Wolf and Koonin, 2013; Albalat and Cañestro, 2016). However, the evidence for genome streamlining obtained with a Birth-Death-Gain model in this analysis was quite modest. This may be due to the flexibility of Birth-Death-Gain models, which do not seek to minimize homoplasy, are agnostic with respect to gains versus losses, and allow for multiple independent origins of the same domain family. In contrast, analysis of the same data with the highly constrained Dollo parsimony model yielded a history dominated by losses. The stark contrast between these inferences prompts a critical reassessment of the methodologies commonly used in ancestral reconstruction.

## Author contributions

Conceptualization and Methodology: DD, MS, LW. Investigation and analysis: YX, MS. Visualization: MS, YX. Writing original draft: DD, MS, YX. Review and editing: DD, LW, MS, YX. All authors read and approved the final manuscript.

## Acknowledgements

The authors thank Dr. Sivaraman Balakrishnan for insightful discussions on data analysis for this work.

## Appendix A Supplementary Materials

### Supplementary Figures

**Fig. A1:**
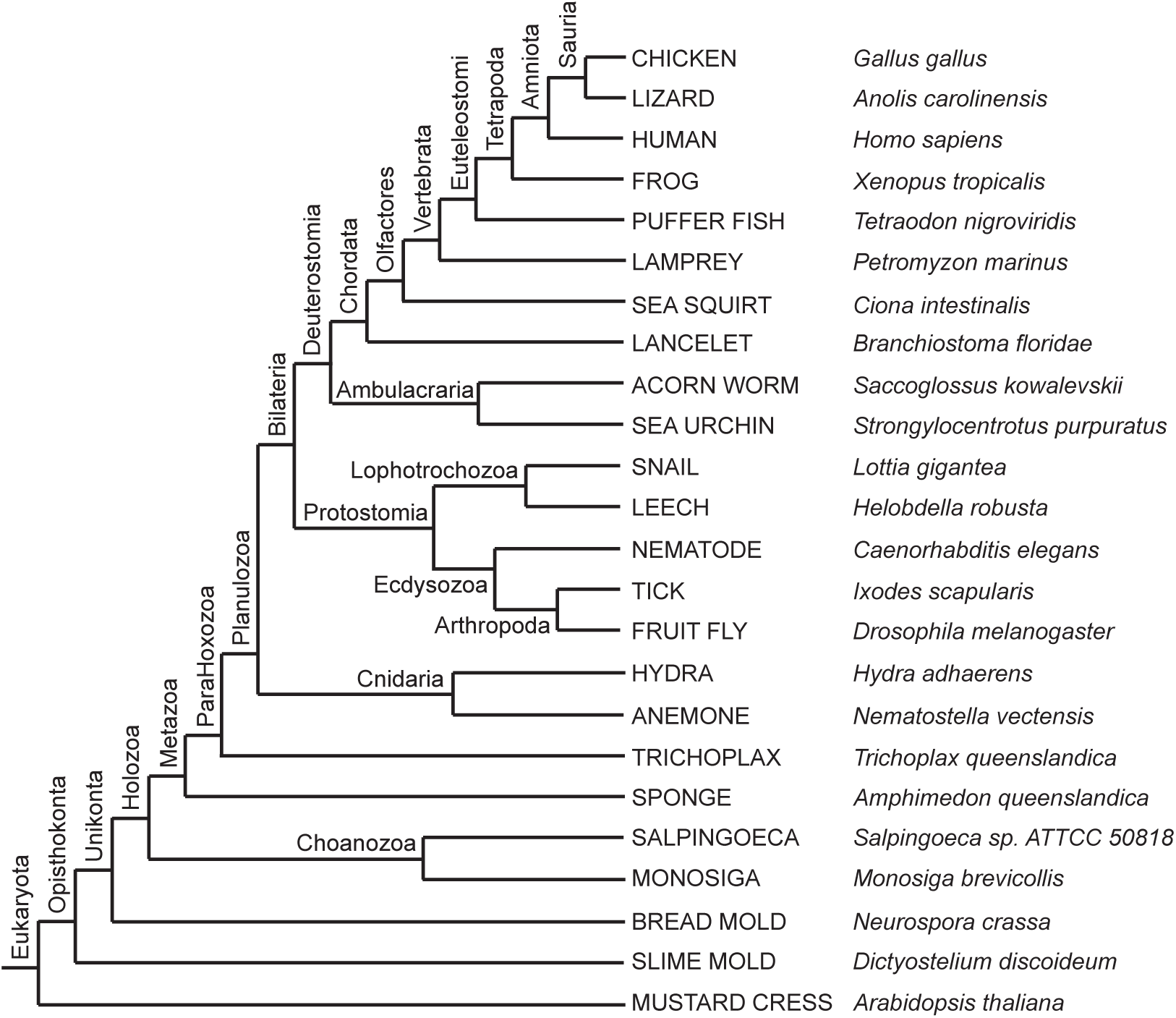
Species tree for all 24 species in the input data, adapted from Philippe et al. (2009).

**Fig. A2:**
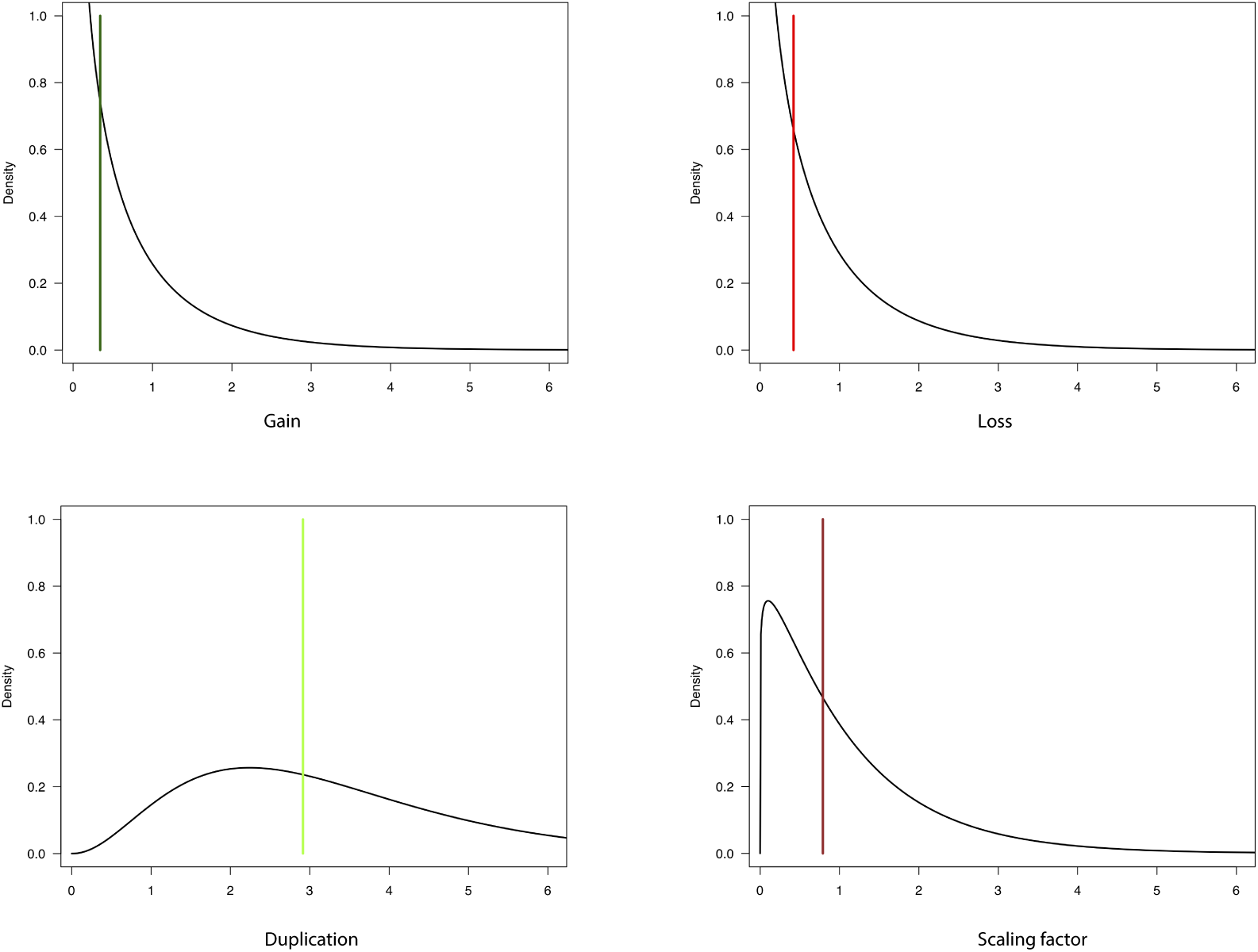
Discretized gamma distribution with two bins for family-specific event rates. Vertical lines indicate the separation between the slow and fast bins.

**Fig. A3:**
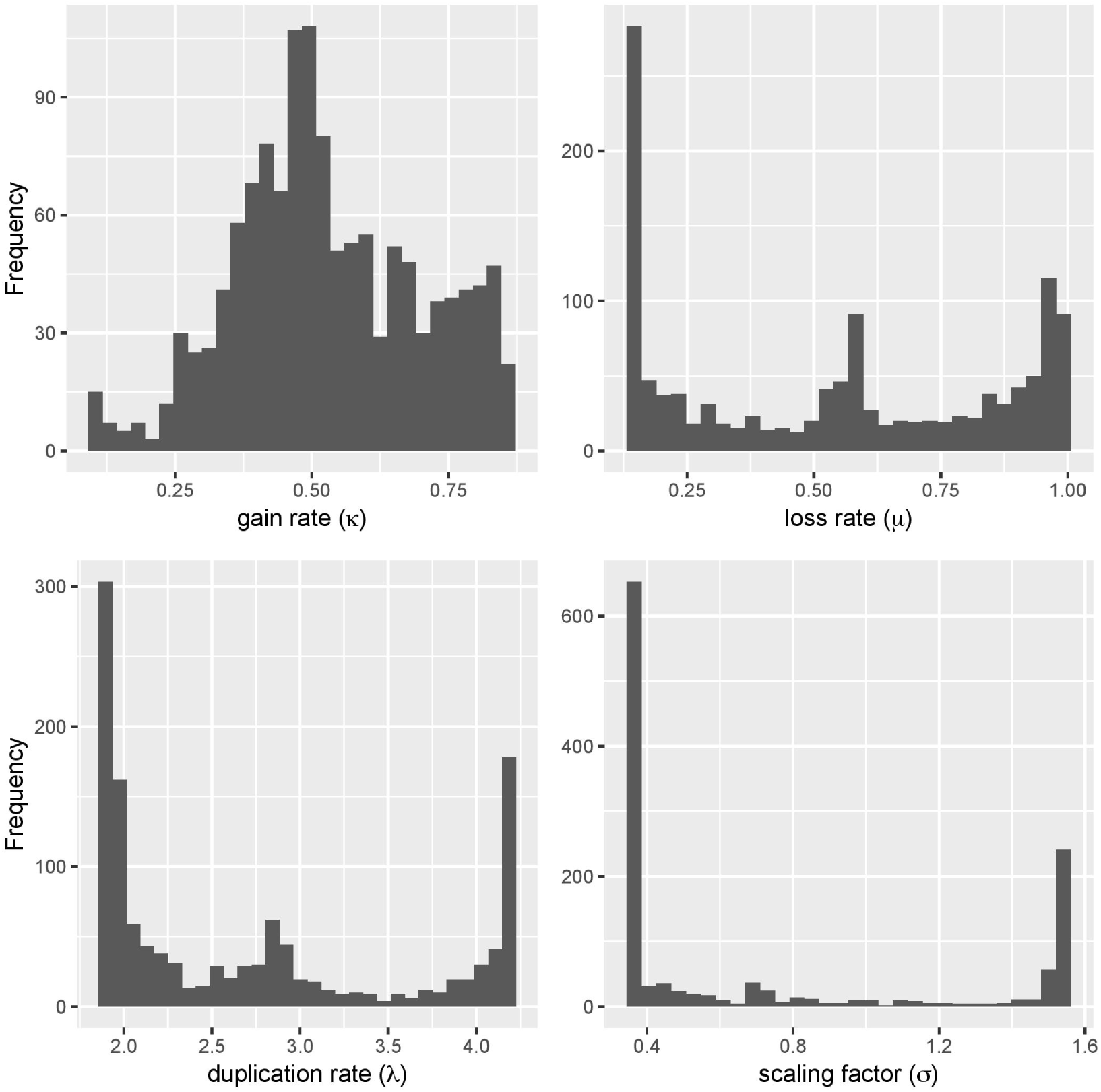
Histograms of expected family-specific rates for the 1283 Holozoan domain families in our data set.

**Fig. A4:**
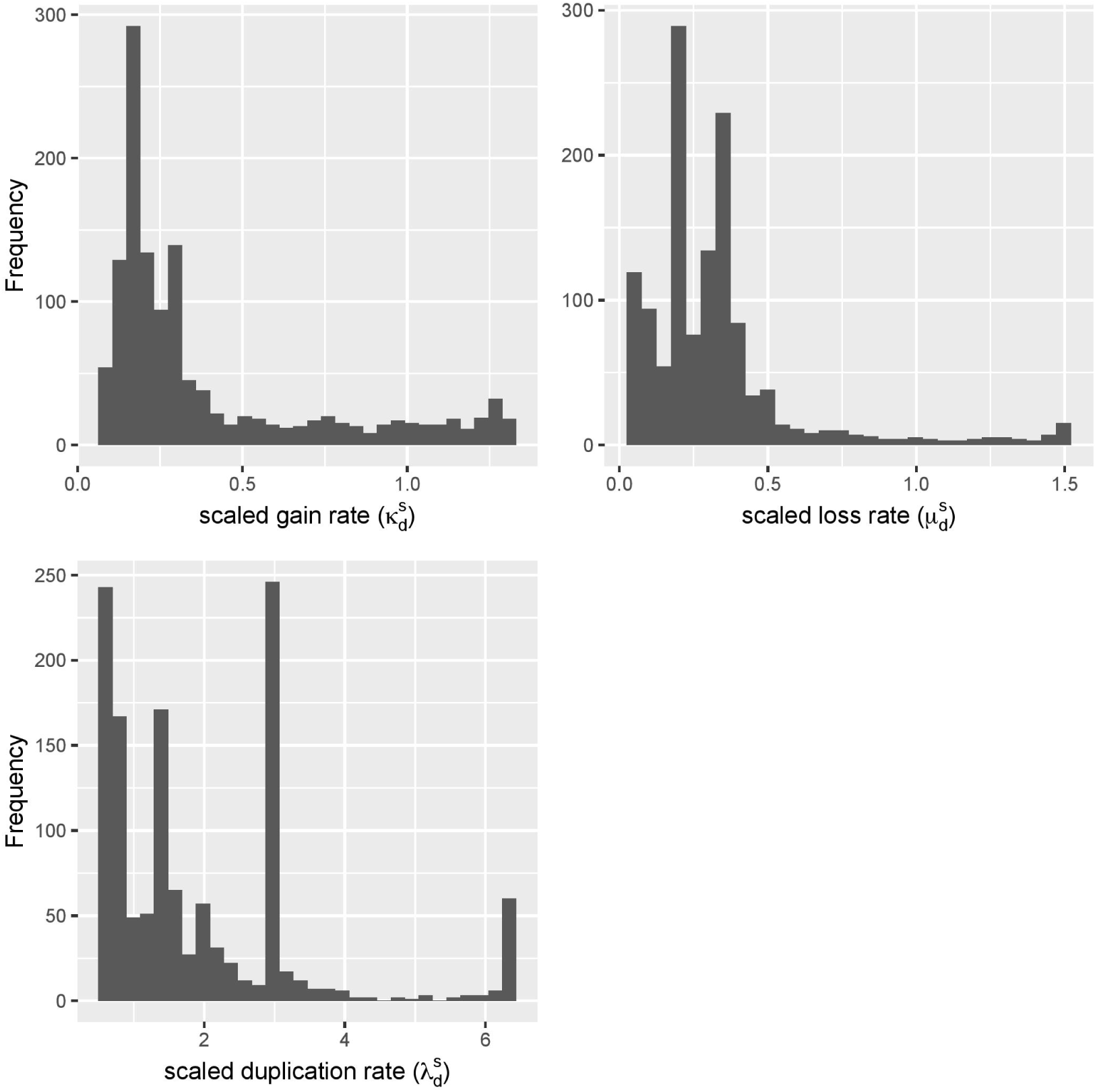
Distributions of expected domain family rates, scaled by scaling factor. The scaled expected rate of each domain family was calculated by multiplying the expected family rate by the expected family scaling factor.

**Fig. A5:**
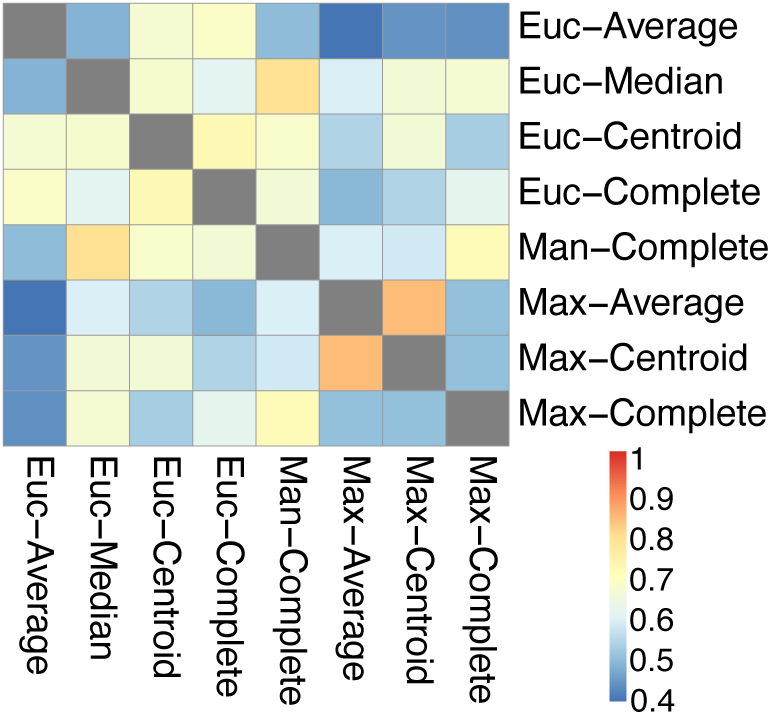
Normalized Mutual Information for all pairs of significant (*p <* 0.05) clusterings with 3 or more clusters. The NMI values range from 0.43 to 1. All diagonal cells are masked in grey.

**Fig. A6:**
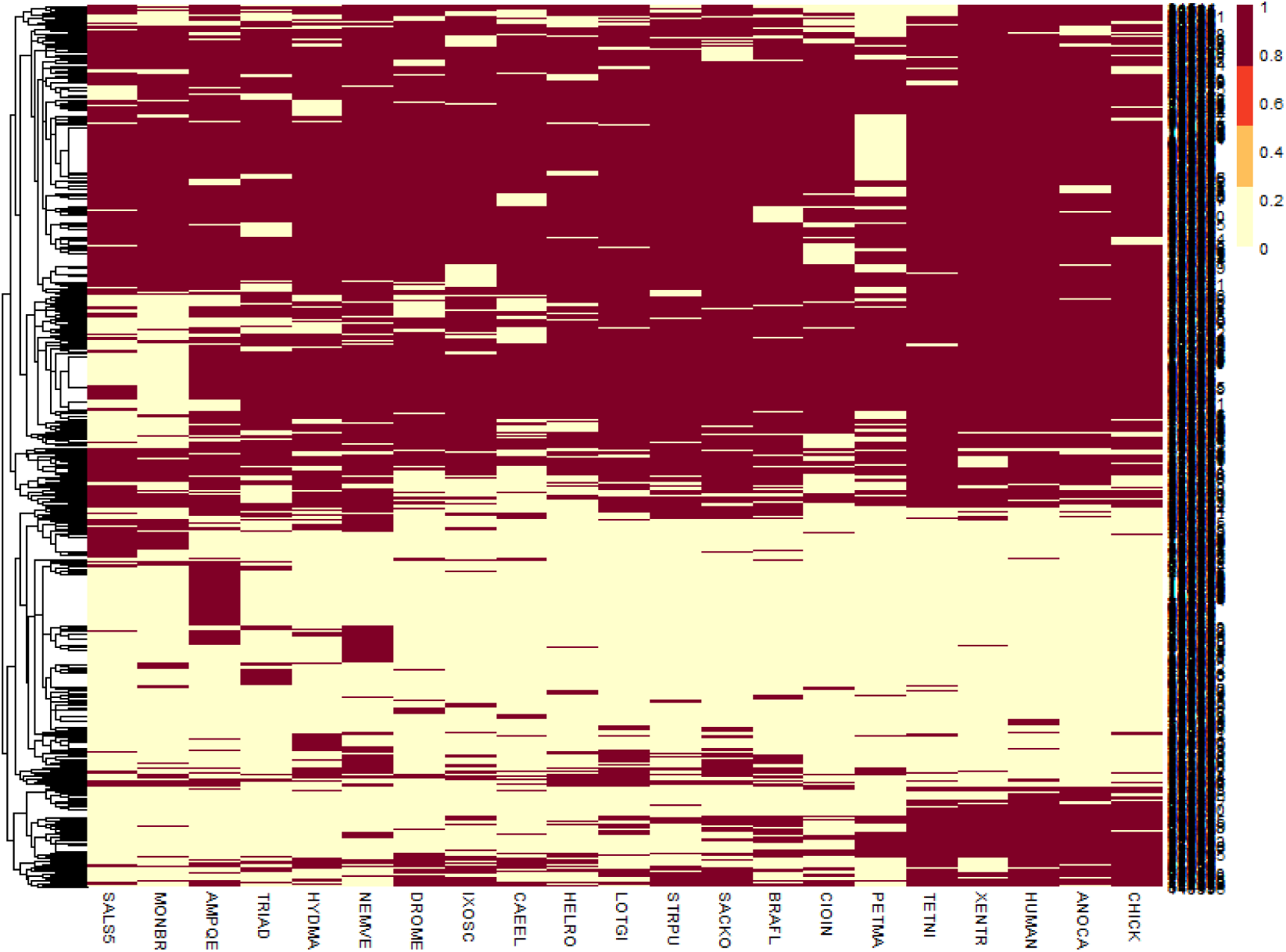
Distribution of the 782 non-universal domain families encoded in 21 holozoan genomes (dark red: present; white: absent). Dendrogram of domain family phylogenetic profiles generated using the pheatmap package in R (Kolde, 2018).

**Fig. A7:**
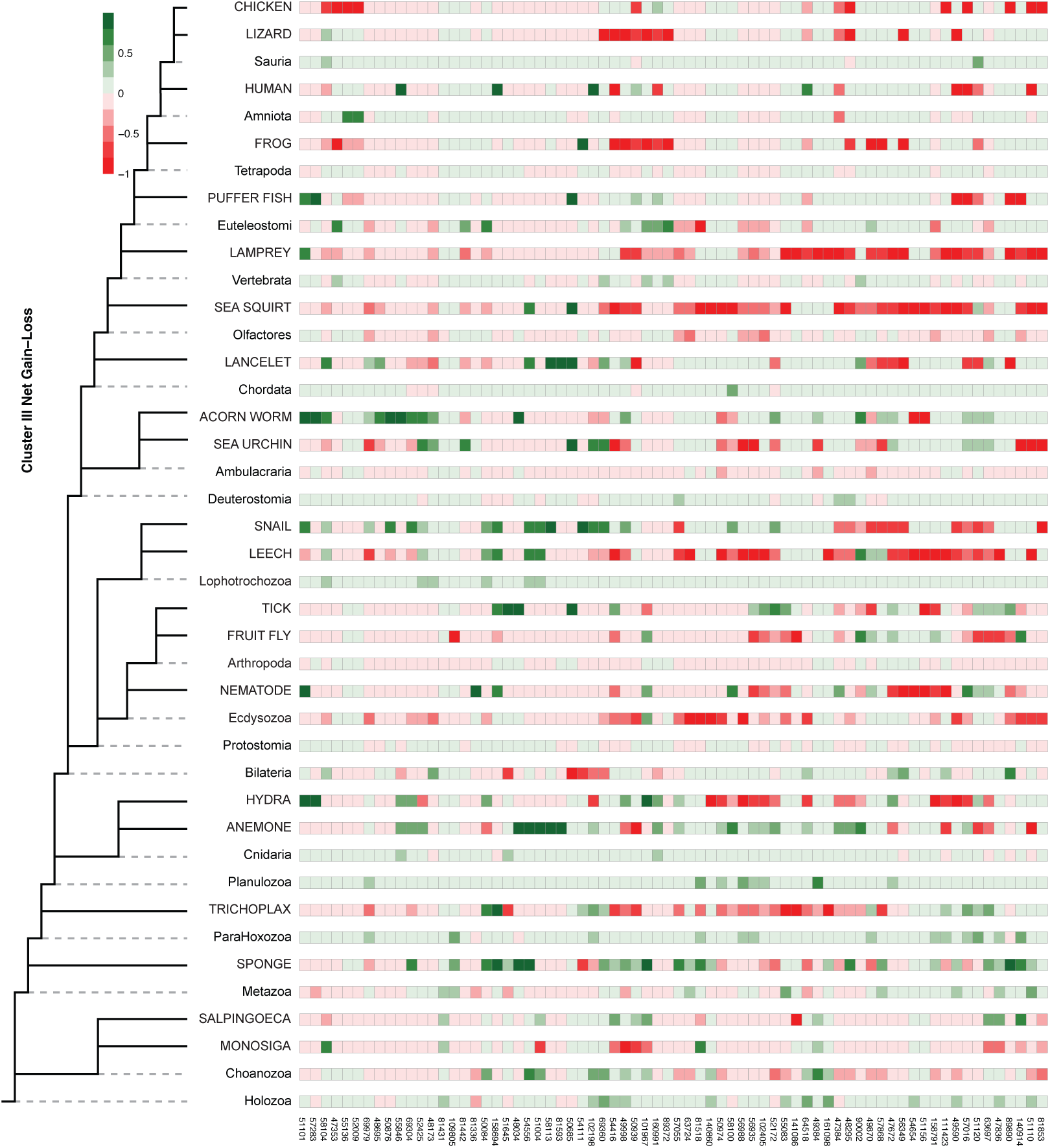
Cluster III gain and loss events.

**Fig. A8:**
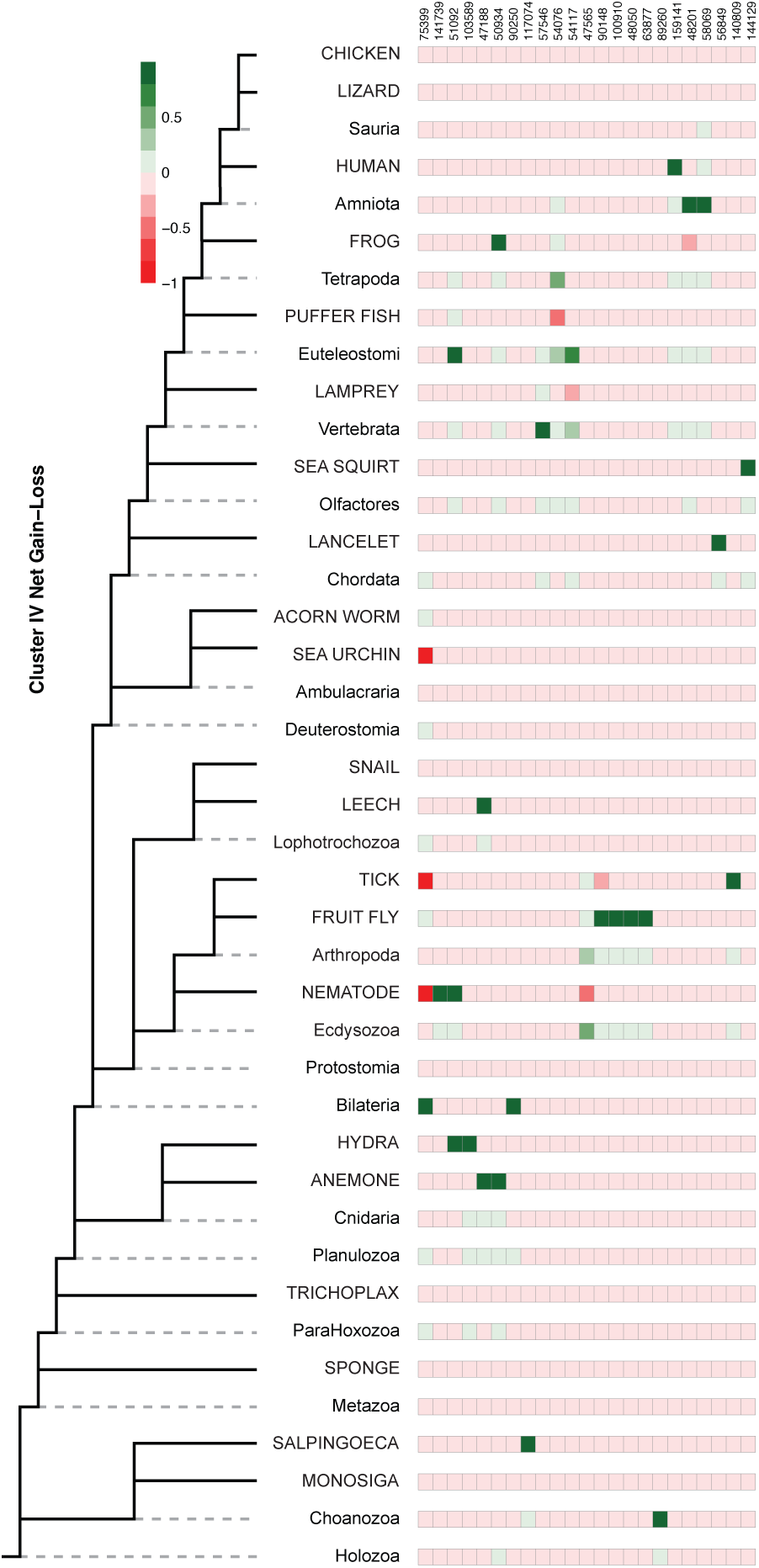
Cluster IV gain and loss events.

**Fig. A9:**
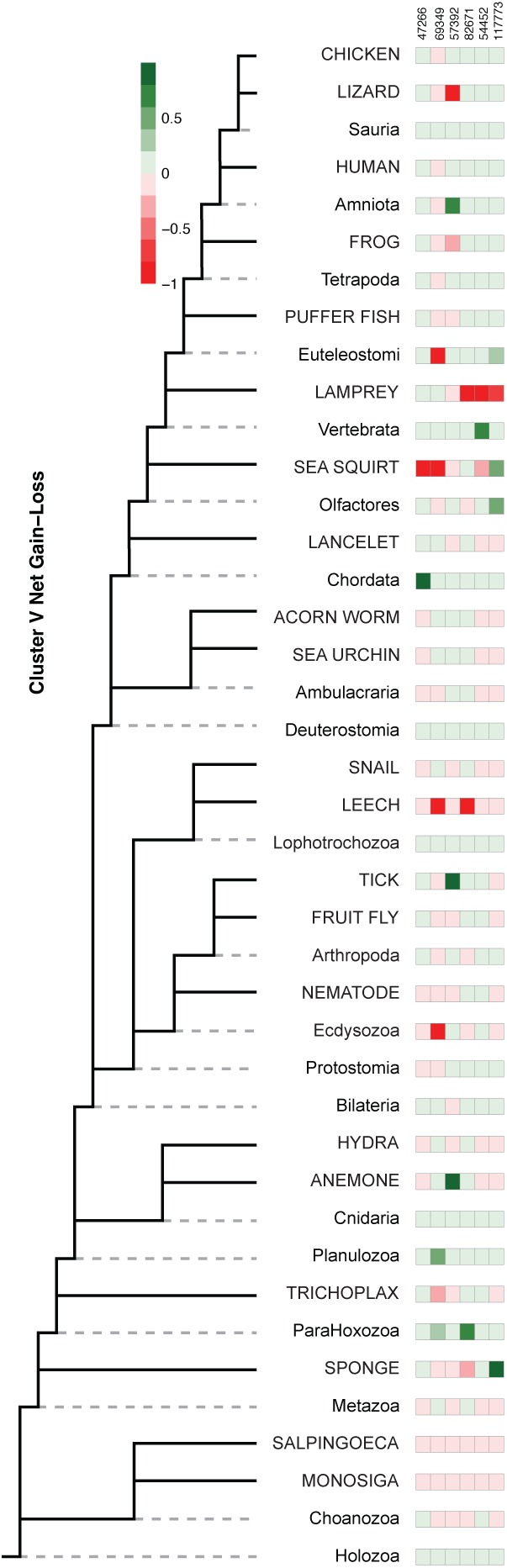
Cluster V gain and loss events.

**Fig. A10:**
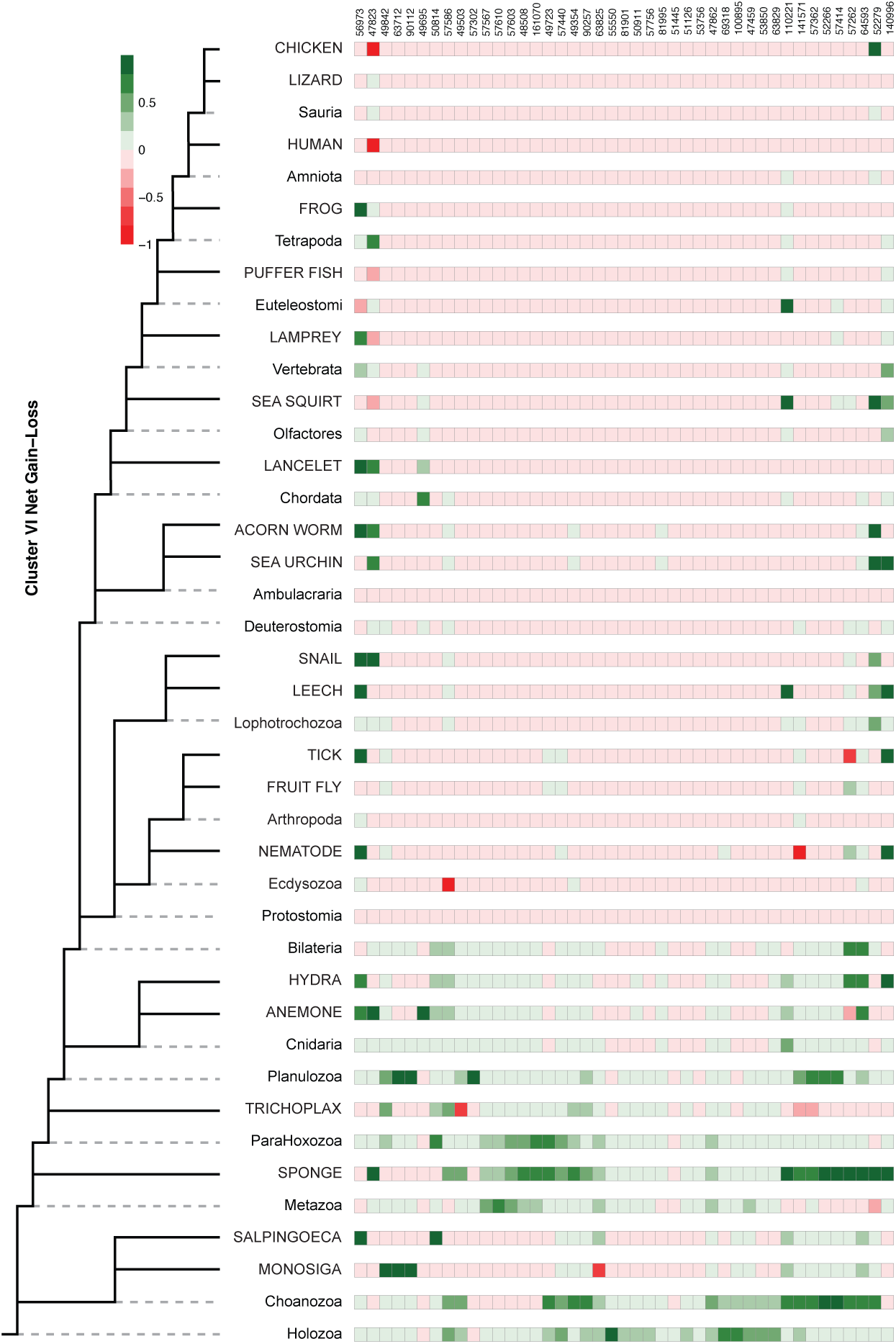
Cluster VI gain and loss events.

### Supplementary Tables

**Table A1:**
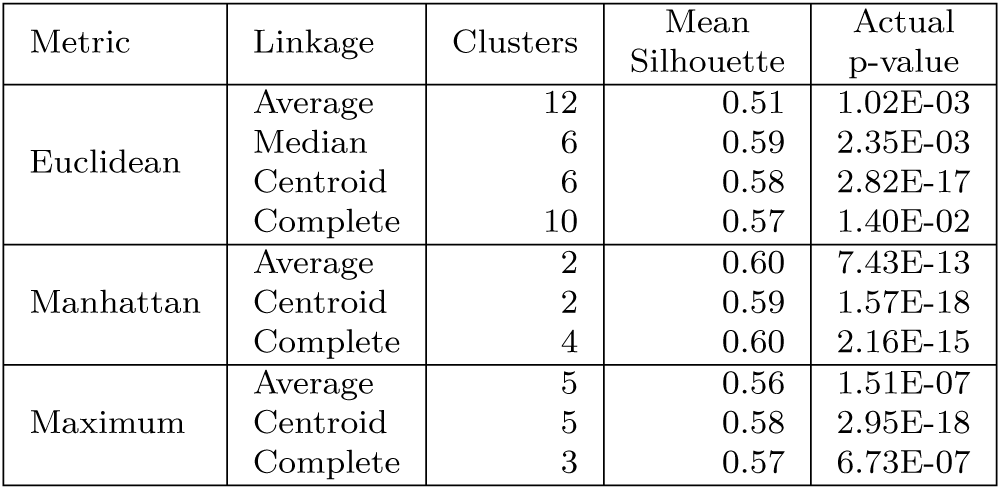
Significant clusterings of scaled event rates (*p <* 0.05).

**Table A2:** Large domain families that were excluded from the Birth-Death-Gain analysis due to convergence problems.

| SUPERFAM ID | Domain Family | Copies in 24 species |  |  |
| --- | --- | --- | --- | --- |
|  |  | Total | Mean | Max |
| 47113 | Histone-fold | 2237 | 93.21 | 385 |
| 47986 | DEATH domain | 2347 | 97.79 | 648 |
| 48264 | Cytochrome P450 | 2396 | 99.83 | 288 |
| 49599 | TRAF domain-like | 1554 | 64.75 | 442 |
| 50494 | Trypsin-like serine proteases | 1888 | 78.67 | 272 |
| 52047 | RNI-like | 2861 | 119.21 | 398 |
| 52200 | Toll/Interleukin receptor TIR domain | 1129 | 47.04 | 293 |
| 53300 | vWA-like | 2527 | 105.29 | 244 |
| 53822 | Periplasmic binding protein-like I | 1499 | 62.46 | 385 |
| 54695 | POZ domain | 3093 | 128.88 | 500 |
| 55486 | Metalloproteases (zincins), catalytic domain | 2607 | 108.62 | 337 |
| 56219 | DNase I-like | 2131 | 88.79 | 620 |
| 56496 | Fibrinogen C-terminal domain-like | 1373 | 57.21 | 271 |
| 56672 | DNA/RNA polymerases | 2521 | 105.04 | 709 |
| 57256 | Elafin-like | 884 | 36.83 | 663 |
| 57625 | Invertebrate chitin-binding proteins | 796 | 33.17 | 284 |
| 57845 | B-box zinc-binding domain | 1865 | 77.71 | 351 |
| 81324 | Voltage-gated potassium channels | 2931 | 122.12 | 283 |
| 101898 | NHL repeat | 1167 | 48.62 | 382 |
| 109640 | KRAB domain (Kruppel-associated box) | 585 | 24.38 | 423 |
| 117281 | Kelch motif | 2077 | 86.54 | 418 |
| 141072 | CalX-like | 1226 | 51.08 | 260 |

**Table A3:**
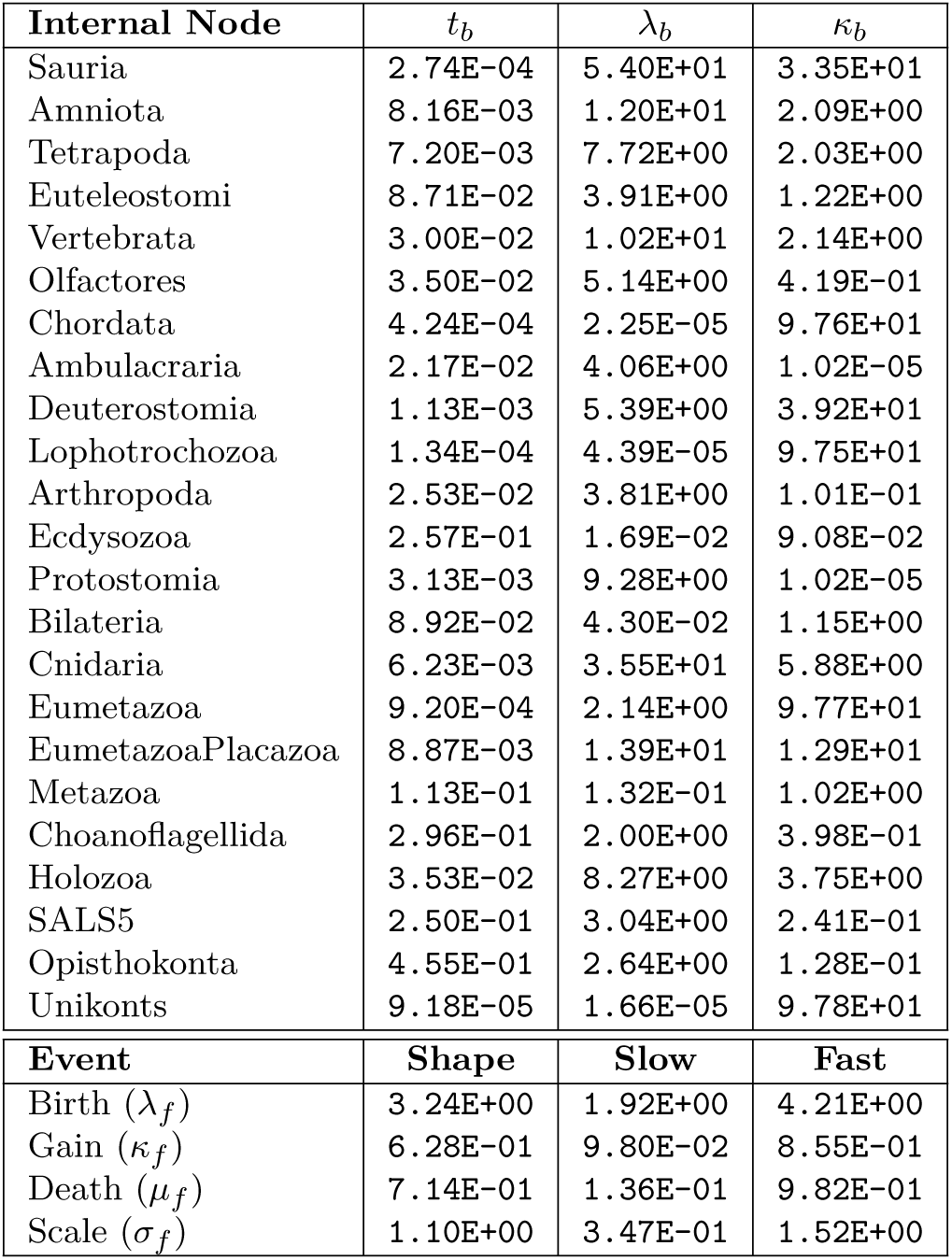
BDG parameters inferred by Count: Branch parameters and midpoints for {*σ_f_, κ_f_, λ_f_, µ_f_* }. Note that loss rates on all internal branches are normalized to be 1.

**Table A4:** Species-specific domain families.

| Species | Copy Numbers |
| --- | --- |
| AMPQE | 37 |
| CAEEL | 4 |
| CIOIN | 1 |
| DROME | 5 |
| HELRO | 2 |
| HUMAN | 6 |
| HYDMA | 3 |
| IXOSC | 3 |
| LOTGI | 1 |
| MONBR | 2 |
| NEMVE | 9 |
| SACKO | 1 |
| SALS5 | 2 |
| TRIAD | 8 |

**Table A5:** Number of universal families in the present-day spaces of each core set and each cluster. Numbers in parentheses are the number of present-day species included in each set.

| Core | Families | Cluster I | Cluster II | Cluster III | Cluster IV | Cluster V | Cluster VI |
| --- | --- | --- | --- | --- | --- | --- | --- |
| All (24) | 445 | 310 | 122 | 1 | 0 | 0 | 12 |
| Holozoan (21) | 501 | 336 | 146 | 1 | 0 | 0 | 18 |
| Metazoan (19) | 547 | 359 | 163 | 1 | 0 | 0 | 24 |
| Bilateria (15) | 594 | 382 | 176 | 1 | 1 | 0 | 34 |
| Euteleost (5) | 899 | 640 | 194 | 16 | 5 | 4 | 40 |

**Table A6:**
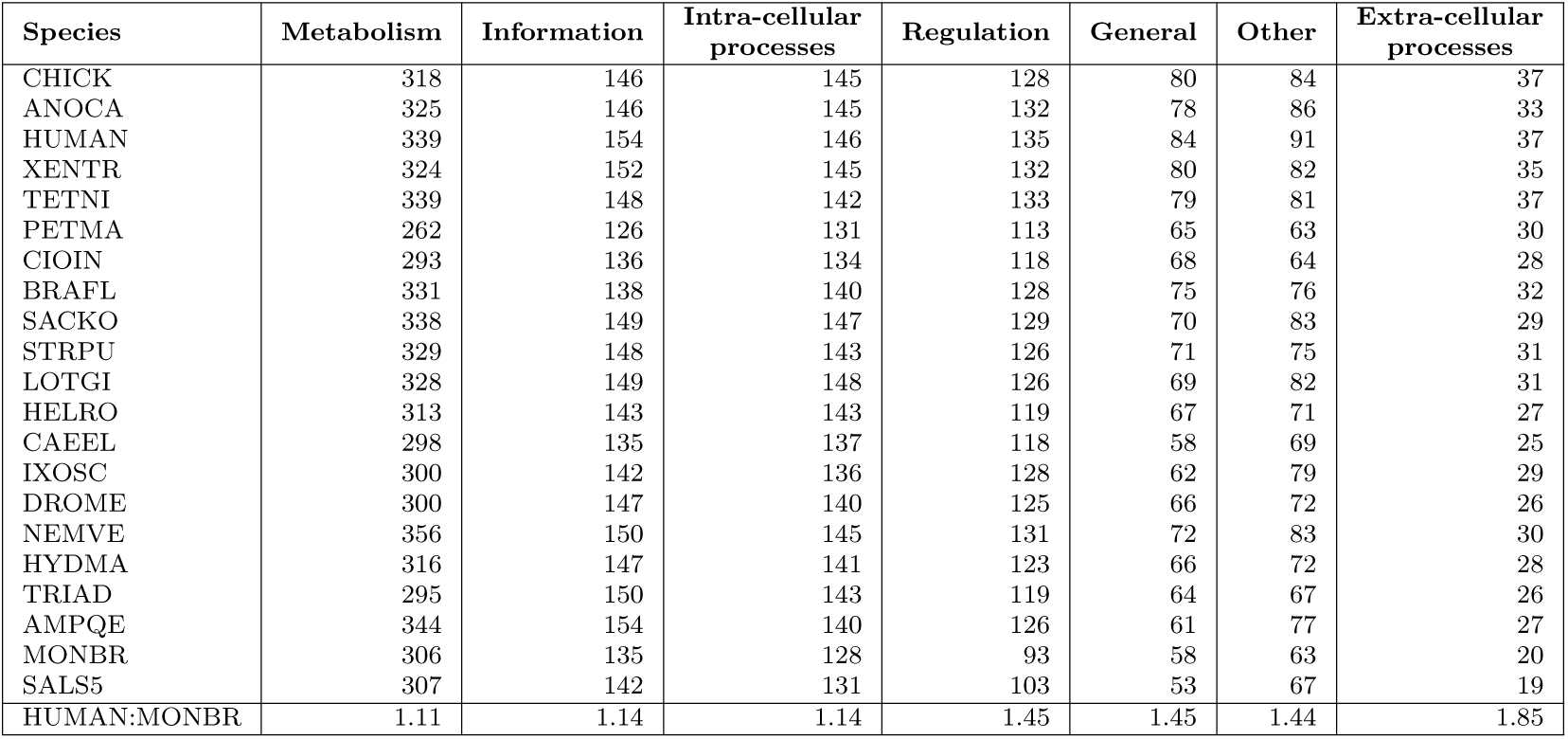
The general functional category distribution across the 1999 holozoan domain families in our data set that have functional annotations.

| Species | Metabolism | Information | Intra-cellular processes | Regulation | General | Other | Extra-cellular processes |
| --- | --- | --- | --- | --- | --- | --- | --- |
| CHICK | 318 | 146 | 145 | 128 | 80 | 84 | 37 |
| ANOCA | 325 | 146 | 145 | 132 | 78 | 86 | 33 |
| HUMAN | 339 | 154 | 146 | 135 | 84 | 91 | 37 |
| XENTR | 324 | 152 | 145 | 132 | 80 | 82 | 35 |
| TETNI | 339 | 148 | 142 | 133 | 79 | 81 | 37 |
| PETMA | 262 | 126 | 131 | 113 | 65 | 63 | 30 |
| CIOIN | 293 | 136 | 134 | 118 | 68 | 64 | 28 |
| BRAFL | 331 | 138 | 140 | 128 | 75 | 76 | 32 |
| SACKO | 338 | 149 | 147 | 129 | 70 | 83 | 29 |
| STRPU | 329 | 148 | 143 | 126 | 71 | 75 | 31 |
| LOTGI | 328 | 149 | 148 | 126 | 69 | 82 | 31 |
| HELRO | 313 | 143 | 143 | 119 | 67 | 71 | 27 |
| CAEEL | 298 | 135 | 137 | 118 | 58 | 69 | 25 |
| IXOSC | 300 | 142 | 136 | 128 | 62 | 79 | 29 |
| DROME | 300 | 147 | 140 | 125 | 66 | 72 | 26 |
| NEMVE | 356 | 150 | 145 | 131 | 72 | 83 | 30 |
| HYDMA | 316 | 147 | 141 | 123 | 66 | 72 | 28 |
| TRIAD | 295 | 150 | 143 | 119 | 64 | 67 | 26 |
| AMPQE | 344 | 154 | 140 | 126 | 61 | 77 | 27 |
| MONBR | 306 | 135 | 128 | 93 | 58 | 63 | 20 |
| SALS5 | 307 | 142 | 131 | 103 | 53 | 67 | 19 |
| HUMAN:MONBR | 1.11 | 1.14 | 1.14 | 1.45 | 1.45 | 1.44 | 1.85 |

**Table A7:** Distribution of Detailed Functions across clusters.

| General | Detailed | I | II | III | IV | V | VI | Total |
| --- | --- | --- | --- | --- | --- | --- | --- | --- |
| Extra-cellular | Cell adhesion | 9 | 9 | 1 | 0 | 1 | 2 | 22 |
|  | Immune response | 1 | 3 | 0 | 1 | 2 | 2 | 9 |
|  | Blood clotting | 0 | 1 | 0 | 0 | 0 | 1 | 2 |
|  | Toxins/defense | 6 | 0 | 3 | 2 | 1 | 1 | 13 |
| General | Small molecule binding | 13 | 2 | 3 | 0 | 0 | 0 | 18 |
|  | Ion binding | 7 | 1 | 1 | 0 | 0 | 0 | 9 |
|  | Lipid/membrane binding | 1 | 0 | 0 | 0 | 0 | 0 | 1 |
|  | Ligand binding | 3 | 0 | 0 | 0 | 0 | 0 | 3 |
|  | General | 14 | 5 | 1 | 0 | 0 | 0 | 20 |
|  | Protein interaction | 28 | 9 | 1 | 1 | 0 | 1 | 40 |
|  | Structural protein | 3 | 0 | 0 | 1 | 0 | 1 | 5 |
| Infor-mation | Chromatin structure | 4 | 1 | 0 | 0 | 0 | 0 | 5 |
|  | Translation | 86 | 2 | 0 | 0 | 0 | 0 | 88 |
|  | Transcription | 21 | 0 | 0 | 0 | 0 | 0 | 21 |
|  | DNA replication/repair | 38 | 6 | 3 | 0 | 0 | 0 | 47 |
|  | RNA processing | 8 | 1 | 0 | 1 | 0 | 0 | 10 |
| Intra-cellular | Cell cycle, Apoptosis | 8 | 6 | 0 | 0 | 0 | 0 | 14 |
|  | Phospholipid m/tr | 2 | 4 | 0 | 0 | 0 | 0 | 6 |
|  | Cell motility | 8 | 6 | 2 | 1 | 0 | 1 | 18 |
|  | Protein modification | 18 | 7 | 0 | 0 | 0 | 3 | 28 |
|  | Proteases | 15 | 15 | 4 | 0 | 0 | 5 | 39 |
|  | Ion m/tr | 12 | 7 | 0 | 0 | 0 | 3 | 22 |
|  | Transport | 34 | 9 | 2 | 0 | 0 | 1 | 46 |
|  | Trafficking/secretion | 2 | 0 | 0 | 0 | 0 | 0 | 2 |
| Metabolism | Energy | 44 | 1 | 5 | 1 | 0 | 0 | 51 |
|  | E- transfer | 15 | 1 | 1 | 1 | 0 | 0 | 18 |
|  | Photosynthesis | 3 | 0 | 0 | 0 | 0 | 0 | 3 |
|  | Amino acids m/tr | 16 | 2 | 0 | 0 | 0 | 0 | 18 |
|  | Nitrogen m/tr | 0 | 1 | 0 | 0 | 0 | 0 | 1 |
|  | Nucleotide m/tr | 21 | 6 | 1 | 0 | 0 | 0 | 28 |
|  | Carbohydrate m/tr | 14 | 2 | 2 | 0 | 0 | 3 | 21 |
|  | Polysaccharide m/tr | 6 | 2 | 4 | 1 | 0 | 2 | 15 |
|  | Coenzyme m/tr | 43 | 3 | 3 | 0 | 0 | 0 | 49 |
|  | Lipid m/tr | 6 | 4 | 1 | 0 | 0 | 2 | 13 |
|  | Cell envelope m/tr | 2 | 0 | 0 | 1 | 0 | 0 | 3 |
|  | Secondary metabolism | 4 | 4 | 1 | 0 | 0 | 0 | 9 |
|  | Redox | 37 | 6 | 4 | 0 | 0 | 0 | 47 |
|  | Transferases | 17 | 6 | 1 | 0 | 0 | 0 | 24 |
|  | Other enzymes | 96 | 19 | 8 | 0 | 0 | 2 | 125 |
| N_A | not annotated | 18 | 0 | 0 | 3 | 0 | 0 | 21 |
| Other | Unknown function | 88 | 5 | 5 | 3 | 0 | 6 | 107 |
|  | Viral proteins | 11 | 0 | 2 | 2 | 1 | 1 | 17 |
| Regulation | RNA binding, m/tr | 19 | 2 | 0 | 0 | 0 | 0 | 21 |
|  | Receptor activity | 5 | 4 | 1 | 1 | 0 | 0 | 11 |
|  | DNA-binding | 37 | 13 | 1 | 0 | 1 | 2 | 54 |
|  | Kinases/phosphatases | 10 | 1 | 1 | 0 | 0 | 0 | 12 |
|  | Signal transduction | 20 | 14 | 3 | 2 | 0 | 3 | 42 |
|  | Other regulatory function | 19 | 2 | 1 | 0 | 0 | 0 | 22 |
| Total |  | 892 | 192 | 66 | 22 | 6 | 42 | 1220 |

**Table A8:** Distribution of General Functions across scaled rate clusters.

| General Function | I | II | III | IV | V | VI | Total |
| --- | --- | --- | --- | --- | --- | --- | --- |
| Extra-cellular processes | 16 (2%) | 13 (7%) | 4 (6%) | 3 (14%) | 4 (67%) | 6 (14%) | 46 (4%) |
| General | 69 (8%) | 17 (9%) | 6 (9%) | 2 (9%) | 0 (0%) | 2 (5%) | 96 (8%) |
| Information | 157 (18%) | 10 (5%) | 3 (5%) | 1 (5%) | 0 (0%) | 0 (0%) | 171 (14%) |
| Intra-cellular processes | 99 (11%) | 54 (28%) | 8 (12%) | 1 (5%) | 0 (0%) | 13 (31%) | 175 (14%) |
| Metabolism | 324 (36%) | 57 (30%) | 31 (47%) | 4 (18%) | 0 (0%) | 9 (21%) | 425 (35%) |
| N_A | 18 (2%) | 0 (0%) | 0 (0%) | 3 (14%) | 0 (0%) | 0 (0%) | 21 (2%) |
| Other | 99 (11%) | 5 (3%) | 7 (11%) | 5 (23%) | 1 (17%) | 7 (17%) | 124 (10%) |
| Regulation | 110 (12%) | 36 (19%) | 7 (11%) | 3 (14%) | 1 (17%) | 5 (12%) | 162 (13%) |
| **Total** | 892 | 192 | 66 | 22 | 6 | 42 | 1220 |

**Table A9:** Significance of Distribution of General Functions.

| General Function | I | II | III | IV - VI |
| --- | --- | --- | --- | --- |
| Extra-cellular processes | **1.98e-06** | 5.96e-01 | 4.04e+00 | **4.60e-06** |
| General | 2.65e+00 | 3.08e+00 | 4.69e+00 | 5.00e+00 |
| Information | **1.22e-10** | **5.27e-05** | 1.72e-01 | **1.23e-02** |
| Intra-cellular processes | **2.02e-05** | **1.08e-05** | 3.47e+00 | 9.23e-01 |
| Metabolism | 6.66e-02 | 6.02e-02 | 4.50e-01 | 8.19e-02 |
| Regulation | 9.52e-01 | 4.50e-01 | 3.34e+00 | 4.08e+00 |
| Viral | 4.53e+00 | 1.12e+00 | 5.05e+00 | 2.47e-01 |

